# Multi-tissue spatial transcriptomics reveals biological age hotspots in mouse and human aging

**DOI:** 10.1101/2025.11.23.689860

**Authors:** Victor Vicente-Alvarez, Cecilia G. de Magalhaes, Dmitrii Glubokov, Adrian Molière, Alexander Tyshkovskiy, Vadim N. Gladyshev

## Abstract

Aging proceeds heterogeneously across tissues, yet how biological age varies within the spatial architecture of individual organs remains poorly understood. Here, we introduce stAge, a framework that quantifies localized transcriptomic age (tAge) from spatial transcriptomics data in mouse and human samples during natural aging and in response to injury, infection, neurodegeneration, and cancer. stAge captures age differences among samples and provides a single multi-tissue model for assessing aging within and across organs. Across tissues and conditions, stAge uncovers robust spatial gradients of biological age and shows that injury and neurodegeneration induce pronounced age acceleration, with stronger responses in older organisms and partial normalization during recovery. With advancing age, tissues develop pronounced hotspots of accelerated aging and coldspots of preserved resilience. Hotspots are enriched for metabolic and immune aging signatures, whereas chromatin-related signatures are associated with coldspots. These findings show that aging is spatially structured within tissues and lay a foundation for developing spatially targeted rejuvenation strategies.

## INTRODUCTION

Aging is associated with an increased risk of chronic diseases and mortality^1^. Understanding its mechanisms, therefore, is crucial for developing interventions to ameliorate late-life morbidity and ultimately extend lifespan. Molecular dynamics across different temporal stages of the aging process have been widely studied and characterized, across animals^2,3^, organs^4,5^, and cells^6^ . However, aging dynamics within the same tissue remain understudied.

Similarly to how different organs age at different rates^4,5^, different regions within tissues may demonstrate heterogeneous aging. Yet this spatial heterogeneity remains largely unexplored across tissues, and methods to systematically analyze spatial aging patterns across multiple tissues are lacking. Spatial transcriptomics (ST) techniques, including fluorescence (fST) and sequencing-based (sST) methods, capture gene expression profiles preserving their spatial context within tissues at near-cellular resolution^7^. While fST typically requires gene imputation for mechanistic insights and sST screens larger gene panels^7,8^, both approaches fundamentally transform our ability to map aging processes. Unlike bulk transcriptomics techniques, which lose the regional variation by averaging across entire tissues, ST reveals how molecular signatures vary spatially. This unique feature enables identification of which tissue microenvironments age faster, how aged cells cluster together, and whether aging propagates through tissues in spatial patterns. This spatial dimension is critical for understanding tissue-level aging dynamics that cannot be captured by traditional approaches.

Quantifying biological age at the molecular level is essential for understanding disease susceptibility, evaluating interventions, and identifying mechanisms that drive age-related decline. Aging clocks are machine learning models trained to predict age or age-related phenotypes from panels of molecular features^9^. Previous efforts have studied aging dynamics using clocks based on fST trained on small gene panels or examining changes in a reduced number of hand-picked age-correlated genes^10^. Adapting multi-tissue transcriptomic aging clocks^11^ for spatial transcriptomics, we study region-specific aging dynamics based on sST data from human and mice tissues under a variety of conditions.

## RESULTS

We developed stAge, a framework to estimate regional biological age from spatial transcriptomic (ST) data by combining metaspot clustering with transcriptomic aging clocks (Fig. 1a–e). For each sample, we apply the SpatialGroup method (constructing a nearest-neighbor graph on spots and applying Leiden clustering), grouping spots into metaspots that are jointly defined by transcriptional similarity and spatial proximity, thereby avoiding reliance on noisy cell-type annotations at single-spot resolution. Because individual ST spots have low coverage and convoluted cell-type composition, we predicted age at the metaspot level rather than per spot.

**Figure 1.**
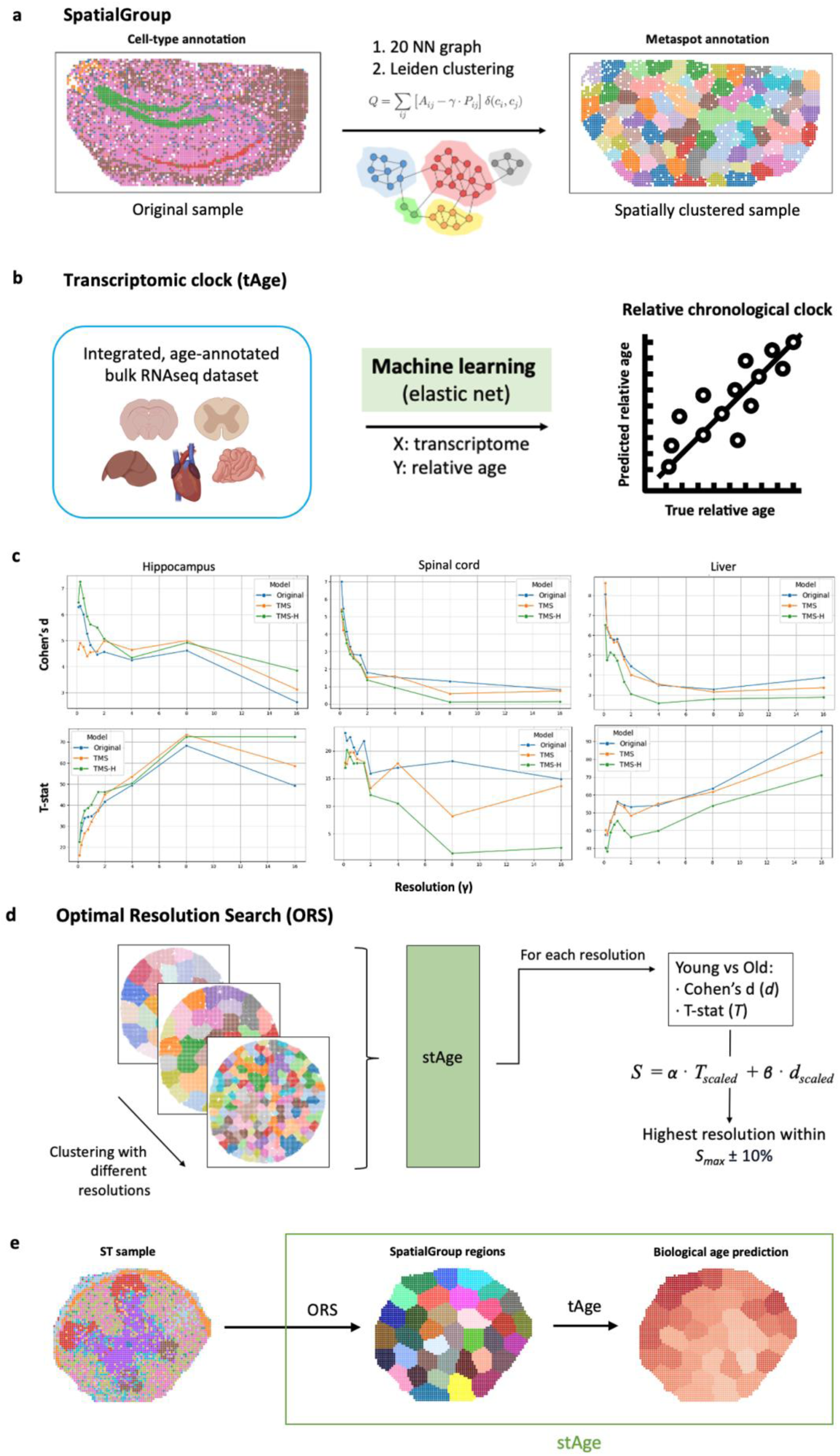
stAge framework. a. SpatialGroup metaspot clustering. Spots are grouped by spatial proximity and transcriptomic similarity by performing Leiden clustering on a spatial connectivity graph. **b. Transcriptomic clock training and validation.** An integrated rodent gene expression meta-dataset with relative chronological age annotation was used to train an elastic net aging multi-tissue clock. The quality of the original rodent multi-tissue transcriptomic clock on test set ^11^ is indicated in the text. **c. Clustering resolution affects size and significance of separation of young and old samples differently across tissues with different clock versions.** The original tAge clock (Original), low coverage-tuned tAge clock (TMS), and high coverage-tuned tAge clock (TMS-H) were used for predictions at different resolution values. Resolution of the Leiden clustering step of the SpatialGroup method determines the size of metaspots and the performance of stAge separating old and young samples across tissues. Increasing resolution achieves more significant separation of young and old metaspots (T-stat increases) but reduces stability of separation (Cohen’s d decreases). **d. Optimal resolution search (ORS).** Due to different performance of transcriptomic clocks across tissues, resolution was maximized for each tissue while ensuring robust age prediction by choosing the highest resolution with maximum composite score (*S_max_*) with 10% tolerance. **e. stAge overview.** For every ST dataset, an optimal SpatialGroup resolution was found via the ORS algorithm, and clustered spatial metaspots were inputted into clocks to generate relative tAge predictions.

We first confirmed that ST data are compatible with our bulk-trained transcriptomic clock. Using the chronological transcriptomic clock, an elastic net machine learning model trained to predict relative age from mouse bulk gene expression data derived from multiple tissues (Fig. 1b), we obtained accurate predictions when applied to fully pseudobulked ST profiles (Fig. S1). Relative chronological transcriptomic age (tAge) predictions (the difference in age between groups using one as the reference, generally young or control) were used because of their higher test-set correlation (r=0.958) compared to absolute age predictions (r=0.938)^11^.

Because metaspot size (and thus coverage) depends on the clustering resolution γ, we systematically evaluated prediction performance across resolutions using the original tAge model. In parallel, we asked whether clocks tuned to lower- or higher-coverage data would further improve performance on ST data. To this end, we generated two additional clock versions (Fig. 1c): TMS, trained by augmenting the bulk training set with low-coverage metacells (bootstrapped single-cell RNA-seq with ∼50,000 counts), and TMS-H, trained with higher-coverage metacells (∼500,000 counts). All three models showed comparable performance on ST data (Fig. S2), and we therefore used the previously validated original tAge clock for all downstream analyses.

To select an appropriate metaspot scale for each tissue, we quantified how well young and old metaspots were separated at each resolution using a composite score *S*(γ) that combines effect size (Cohen’s d) and statistical significance (t-statistic) (Fig. 1d). Across tissues, increasing resolution generally improved statistical separation of young and old metaspots (higher t-statistic) but reduced effect size (lower Cohen’s d), indicating a trade-off between granularity and robustness. For each tissue, we therefore defined an optimal resolution as any γ within *S_max_* ± 10%, yielding the smallest metaspots that still maintained robust discrimination between age groups. Together, optimal resolution search, SpatialGroup clustering, and tAge prediction constitute the stAge framework (Fig. 1e).

### stAge reveals regional gradients of aging across multiple mouse tissues

We next applied stAge to 10 datasets^12–16^ spanning 8 mouse tissues and visualized biological age predictions for metaspots (Figs. 2, 3). The multi-tissue transcriptomic clock robustly captured chronological age difference of ST samples across tissues, including brain, hippocampus, spinal cord, spleen, liver, intestine, testis, and ovaries (Figs. S3, S4).

**Figure 2.**
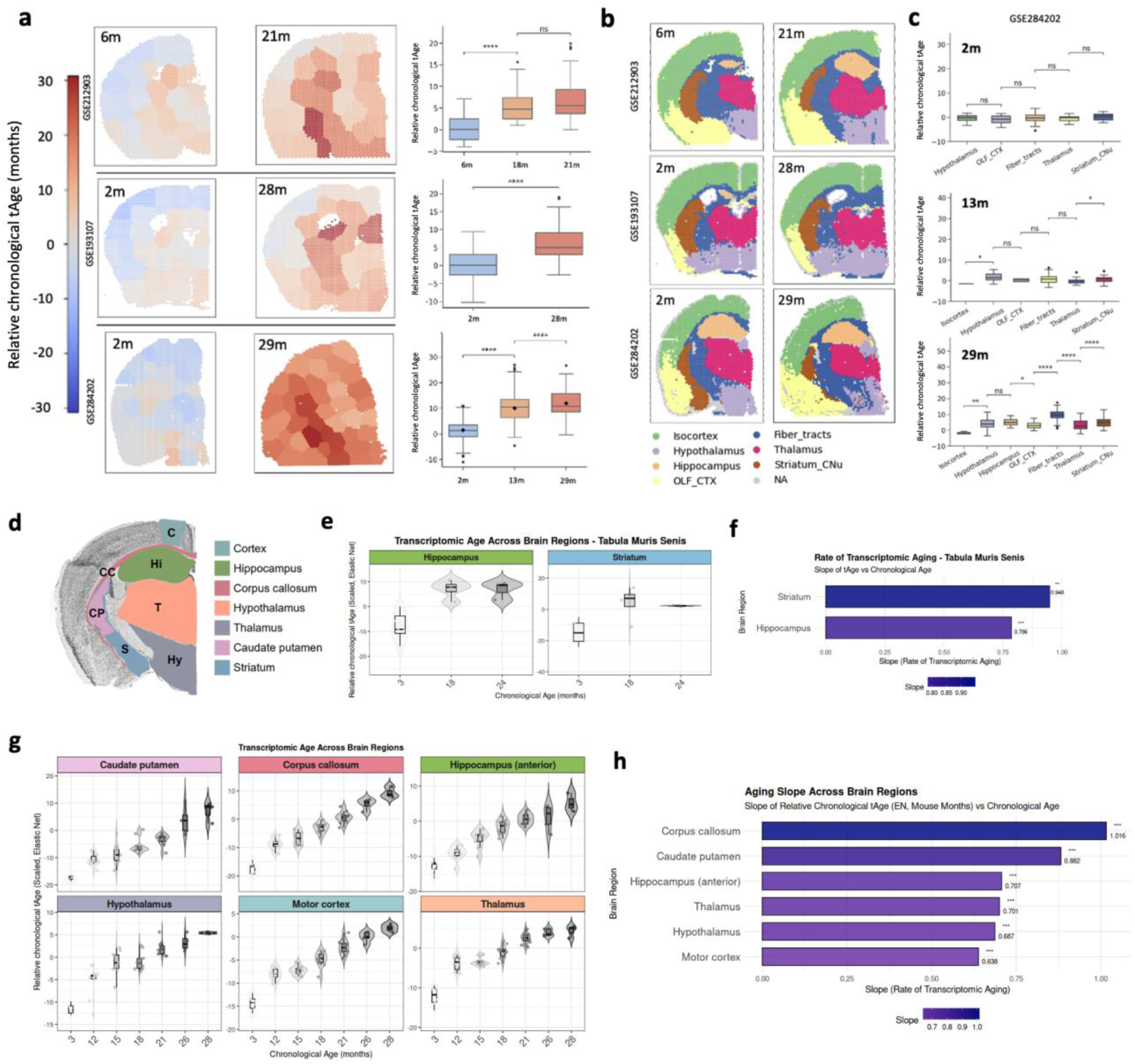
stAge captures local and global age difference in the mouse brain. a. Spatial age difference in the brain. Spatial and box plots of stAge predictions (in months) in three separate brain datasets. One young and one old representative sample in each dataset were selected for spatial plots. Given names for dataset are brain25 (GSE284202), brain2g (GSE193107), and brain3g (GSE212903). In box plots, all samples’ metaspots are displayed for each time point. Mann-Whitney U test. **b. Brain region annotation.** Brain regions were annotated using well-established gene markers into 7 regions. Mann-Whitney U test. Significance: *P* < 0.05 (*), *P* < 0.01 (**), *P* < 0.001 (***), *P <* 0.0001 (****); adjusted P-values shown. **c. Transcriptomic age across brain regions.** Age predictions were generated and displayed in box plots at the three different time points of the GSE284202 dataset. **d. Schematic of brain regions used for single-cell validation.** tAge calculations using publicly available datasets (panels e–h). Brain illustration adapted from the Allen Brain Atlas (https://atlas.brain-map.org). **e. Transcriptomic age of hippocampus and striatum (fiber tracts) in single-cell data from Tabula Muris Senis.** tAge trajectories calculated from single-nucleus RNA-seq data (GSE132042), showing region-specific transcriptomic aging patterns. **f. Slope of transcriptomic age as a function of age in hippocampus and striatum.** Regional tAge progression rates from panel *e*. The striatum, which encompasses the caudate putamen region, displays higher slopes than the hippocampus, consistent with bulk RNA-seq findings. **g. Transcriptomic age of bulk data from six different brain regions.** tAge trajectories across chronological age calculated from bulk RNA-seq data (GSE212336). All analyzed brain regions show progressive increases in transcriptomic age with chronological age. **h. Slope of transcriptomic age as a function of age in six different brain regions.** Regional variation in tAge progression rates from panel *g*.

In the mouse brain, we analyzed three independent datasets (Fig. 2a) and stAge was able to capture age difference between the different age groups. Brain regions were annotated using established marker genes (Fig. 2b)^13^ and inputted to stAge separately. Longitudinal analysis across three ages (young, middle, old; Fig. 2c) showed relatively uniform tAge in young mice (2 months), followed by the emergence of focally accelerated aging in white-matter–rich fiber tracts in old mice (29 months). These observations are consistent with previous reports linking white matter to heightened glial aging and inflammation^13,16^.

To validate these regional patterns, we analyzed bulk RNA-seq and snRNA-seq datasets from dissected mouse brain regions aligned with the coronal plane used in ST (Fig. 2d–h). In agreement with the spatial analysis, white matter tracts (corpus callosum and striatum) showed steeper tAge slopes with chronological age than other regions, confirming accelerated transcriptomic aging in these areas at both spatial and bulk/ single-nucleus resolutions.

stAge also captured age differences in other tissues. In hippocampus, spinal cord, spleen, liver, testis and ovary, metaspots from old samples had significantly higher tAge than those from young samples (Fig. 3a, Fig. S3–S4). The small intestine provided a particularly clear example of spatial age polarity (Fig. 3b). After classifying metaspots into high-turnover (enterocytes, goblet cells, stem/crypt cells) and low-turnover (lymphocytes, Paneth cells, plasmocytes, smooth muscle) categories (Fig. 3c), high-turnover regions consistently showed lower tAge than low-turnover regions (Fig. 3d), consistent with rapid cell renewal and proliferative gene programs buffering aging signatures.

**Figure 3.**
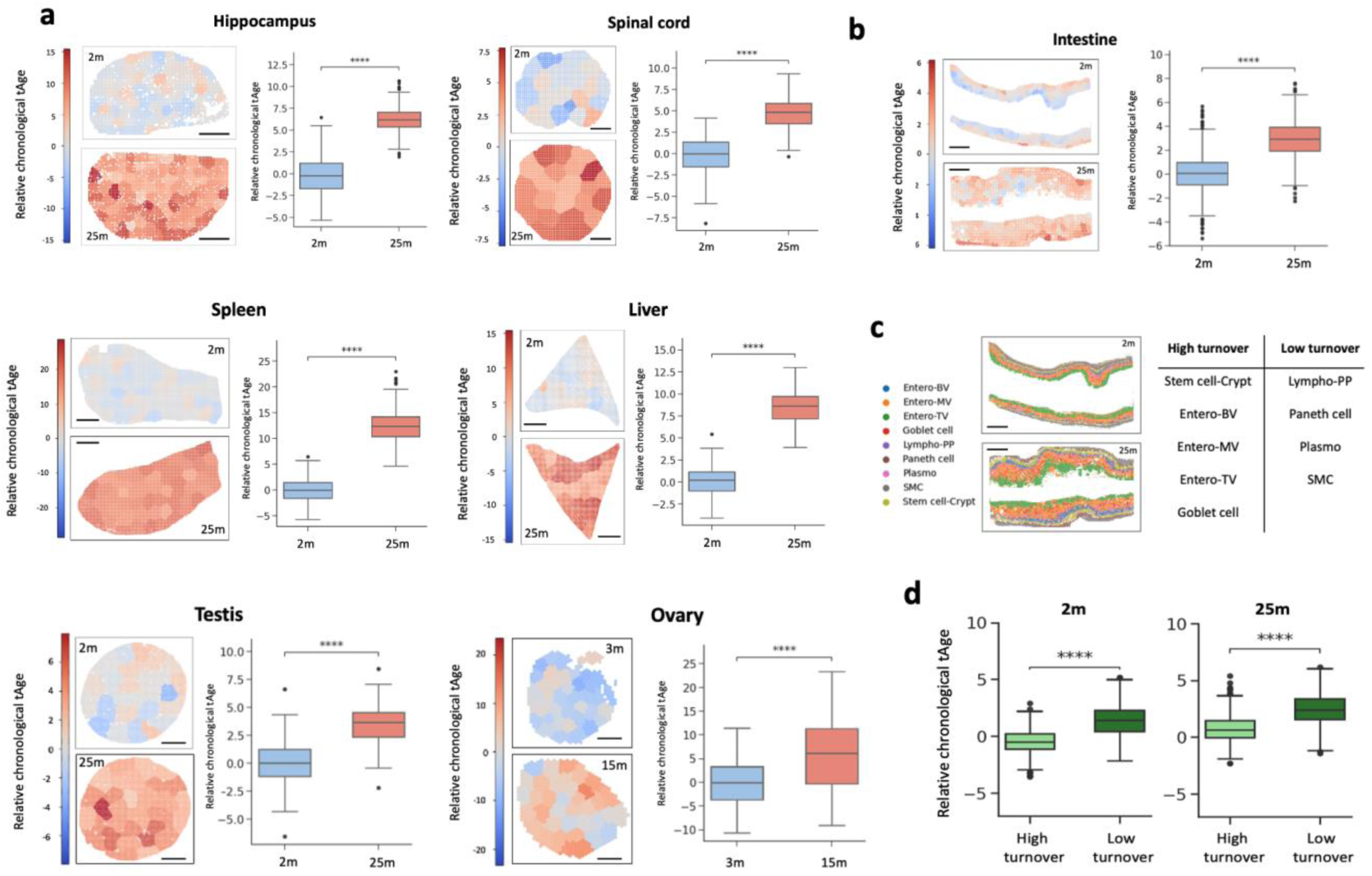
stAge captures age difference in multiple tissues. a. Spatial age difference across multiple tissues. Spatial and box plots of stAge predictions (in months) in six different mouse tissues: Hippocampus, Spinal cord, Spleen, Liver, Testis (STT0000039) and Ovary (GSE188257). One young and one old representative sample in each dataset were selected for spatial plots. In box plots, all samples’ metaspots are displayed for each time point. Scale bars: 500 μm (Hippocampus and Spinal cord), 1 mm others. Mann-Whitney U test. **b. Spatial age difference in the intestine.** Spatial and box plots of stAge predictions in the small intestine dataset. Scale bars 1 mm. Mann-Whitney U test. **c. Intestine spot type architecture and classification.** Spot types provided by the authors include stem and crypt cells (Stem cell-Crypt), enterocytes (Entero-TV, Entero-MV, Entero-BV), lymphoid cells (Lympho-PP), plasmocytes (Plasmo), smooth muscle cells (SMC), Paneth cells, and Goblet cells. Spot types were separated into high turnover and low turnover proliferation cell types. **d. Transcriptomic age of intestinal regions.** Relative tAge predictions of metaspots in high-turnover and low-turnover regions were compared. Mann-Whitney U test. Significance: *P* < 0.05 (*), *P* < 0.01 (**), *P* < 0.001 (***), *P <* 0.0001 (****); adjusted P-values shown.

### Damage and regeneration induce localized age acceleration captured by stAge

We next asked whether stAge captures biological age changes induced by tissue injury and regeneration (Fig. 4). In three independent age-matched intervention datasets (spinal cord crush injury^17,18^, myocardial infarction^18^, bone fracture^19^), injured tissues exhibited markedly higher tAge than sham controls, with partial recovery during regeneration.

**Figure 4.**
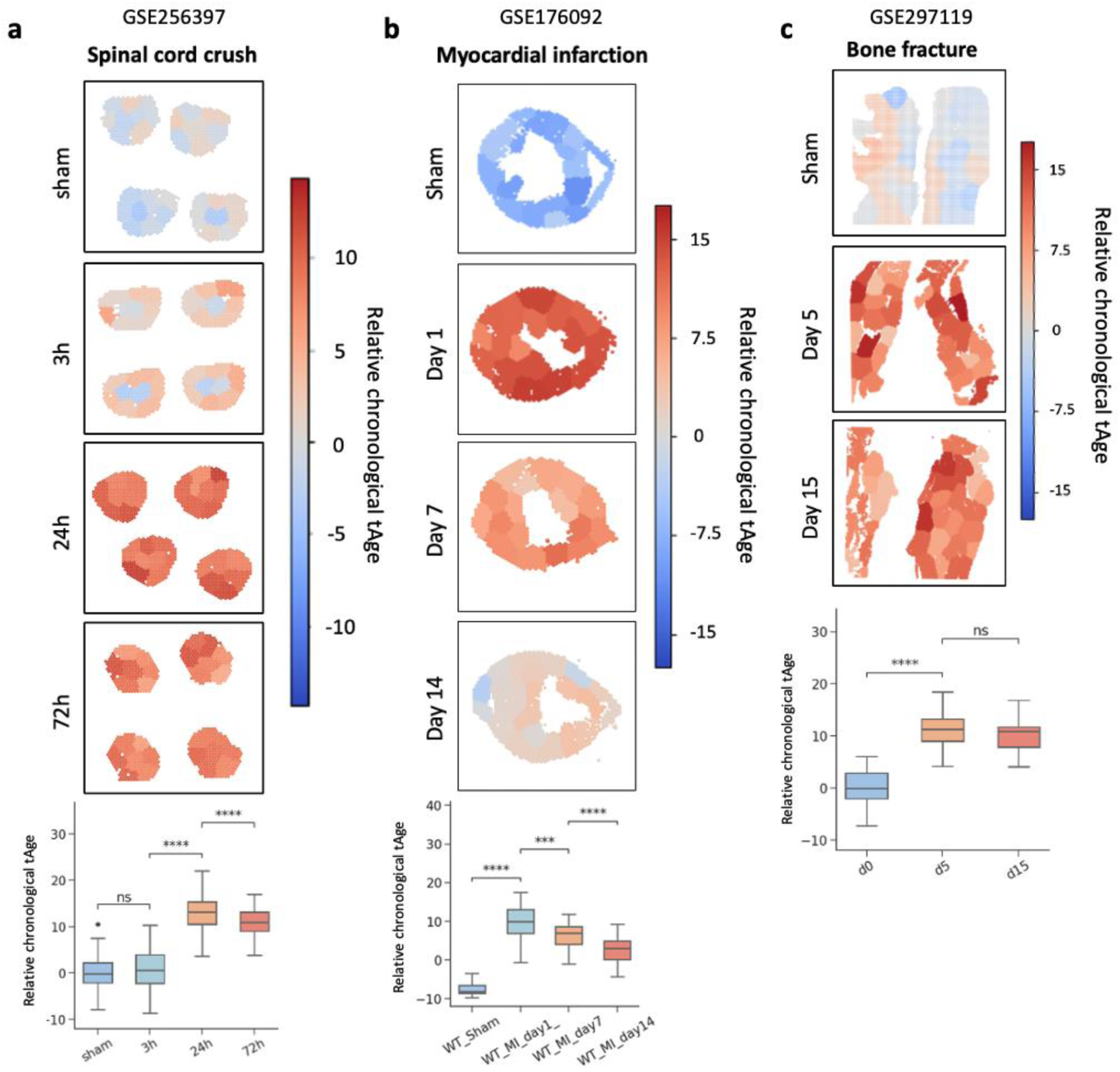
stAge captures tissue damage and recovery. a. Spatial transcriptomic age predictions during spinal cord crush injury progression. Spatial plots display stAge predictions (in months) in age-matched sham and injured mice at 3h, 24h, and 72h after intervention. Box plots display stAge predictions across all samples in each category group (Mann-Whitney U test). **b. Spatial transcriptomic age predictions during myocardial infarction progression.** Spatial plots display stAge predictions in age-matched sham and injured mice at 1 day, 7 days, and 14 days after intervention. Box plots display stAge predictions across all samples in each category group (Mann-Whitney U test). **b. Spatial transcriptomic age predictions during bone fracture progression.** Spatial plots display stAge predictions in age-matched sham and injured mice at 5 days, and 15 days after intervention. Box plots display stAge predictions across all samples in each category group. Mann-Whitney U test. Significance: *P* < 0.05 (*), *P* < 0.01 (**), *P* < 0.001 (***), *P <* 0.0001 (****); adjusted P-values shown.

In the spinal cord crush dataset (Fig. 4a), sham samples displayed the lowest tAge, while post-injury tissues showed progressive age acceleration. White matter regions at the periphery of the cord, enriched for myelinated axonal tracts, already showed slightly elevated tAge 3 h after injury. By 24 h, tAge was strongly increased across nearly the entire spinal cord, coinciding with widespread necrosis. Partial recovery at 72 h reduced but did not normalize tAge, indicating incomplete restoration of transcriptomic age.

A similar pattern was observed in myocardial infarction (Fig. 4b). Sham hearts had low tAge, which rose sharply in infarcted regions, then declined as tissue regenerated but remained above baseline even at later time points. In the bone fracture model (Fig. 4c), trauma induced a strong tAge increase; regeneration proceeded more slowly than in other tissues, and tAge remained elevated even after 15 days, indicating persistent molecular damage. Notably, in all three models, localized lesions were accompanied by age acceleration extending beyond the directly injured areas, suggesting broader tissue remodeling.

Together, these results show that injury and regeneration across multiple tissues are accompanied by rapid, spatially structured increases in transcriptomic age, consistent with the idea that molecular damage signatures that accumulate during aging are acutely engaged during tissue injury and disease.

### Older tissues undergo increased injury- and disease-mediated age acceleration

We then investigated how chronological age modulates the response of tissues to interventions using ST datasets with treatments across age groups^20–22^ (Fig. 5). Across conditions, younger mice had lower overall tAge and showed attenuated treatment-mediated age acceleration, whereas older mice displayed higher baseline tAge and larger treatment-induced increases.

**Figure 5.**
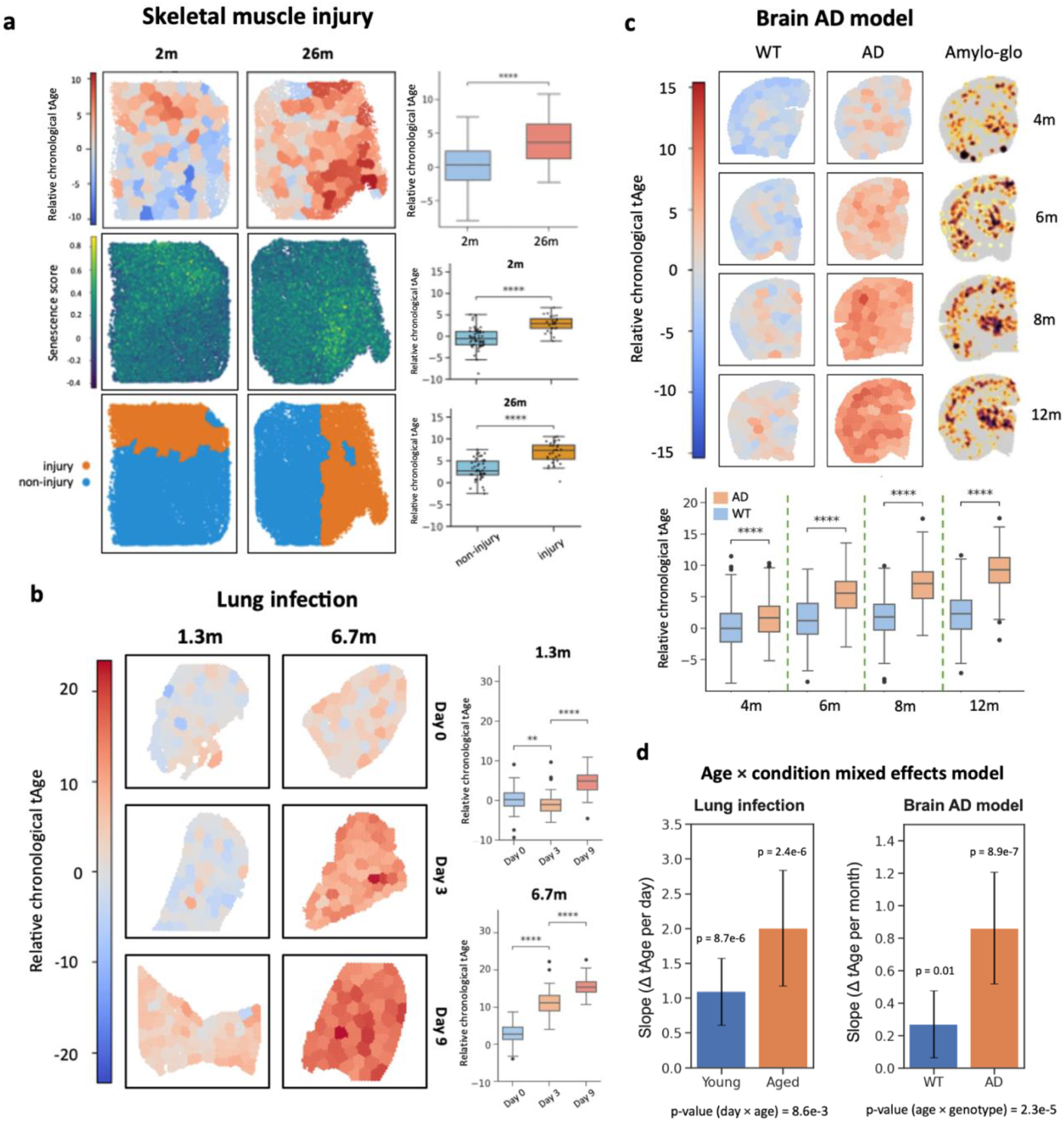
Older tissues undergo increased injury- and disease-mediated age acceleration. a. Spatial age difference in skeletal muscle injury. Spatial plots display stAge predictions (in months), senescence score, and injury site on myotoxin injury in skeletal muscle. (GSE266933) Box plots display stAge prediction differences between young (2m) and old (26m) between young and old samples, and between injury and non-injury sites in young and in old. Mann-Whitney U test. **b. Spatial age difference during lung infection.** Spatial plots display stAge predictions in young (1.3m) and aged (6.7) mice in days 0, 3, and 9 of infection (GSE202322). Box plots display stAge predictions in young and aged mice (Mann-Whitney U test). **c. Spatial age difference during Alzheimer’s disease mouse model lifespan.** Spatial plots display stAge predictions of WT and 5X mouse AD models at 2, 4, 6, 8, and 12 months (GSE233208). Box plots display metaspot stAge predictions in WT and 5X mice at 2, 4, 6, 8, and 12 months (Mann-Whitney U test). Amylo-glo images adapted from Miyoshi et al., (2024)^22^. Significance: *P* < 0.05 (*), *P* < 0.01 (**), *P* < 0.001 (***), *P <* 0.0001 (****); adjusted P-values shown. **d. Mixed effects modeling reveals the additive effect of aging and disease in biological age acceleration.** Significant difference in mixed effects model slopes shows higher age acceleration in aged mice during infection and higher age acceleration during aging of the AD model brain.

In a myotoxin-induced muscle injury model (Fig. 5a), old muscle had significantly higher baseline tAge than young muscle. Within each age group, injury regions showed substantially elevated tAge compared to unaffected muscle, and these regions coincided with high senescence scores, indicating local accumulation of senescence-associated transcriptional programs. However, age acceleration in response to injury was consistently greater in old muscle, suggesting diminished buffering capacity with age.

In an infection-induced lung disease model (Fig. 5b), tAge increased progressively with infection severity. Age acceleration was again stronger in old than in young lungs, highlighting an additive effect of chronological age and disease burden.

Chronic neurodegeneration provided a complementary setting to study treatment-mediated age acceleration. In the 5xFAD model of Alzheimer’s disease (Fig. 5c), wild-type brains exhibited the expected pattern of higher tAge around the fiber tracts, whereas 5xFAD brains showed widespread transcriptomic age acceleration across the entire section, with the strongest tAge increases in cortical regions corresponding to dense amyloid-β plaque deposition. Mixed-effects models across intervention datasets (lung infection–fibrosis and 5xFAD) confirmed a steeper increase in intervention-mediated tAge in older or diseased animals (Fig. 5d), indicating that aging and pathology synergistically amplify regional age acceleration.

### Spatial signatures of biological age hotspots emerge and converge with age

To dissect intra-tissue heterogeneity in biological age, we applied Getis–Ord hotspot analysis^23^ to stAge predictions, identifying regions with significantly higher tAge (hotspots) or lower tAge (coldspots) within each sample (Fig. 6a). We then asked what transcriptional and cellular features distinguish these regions.

**Figure 6.**
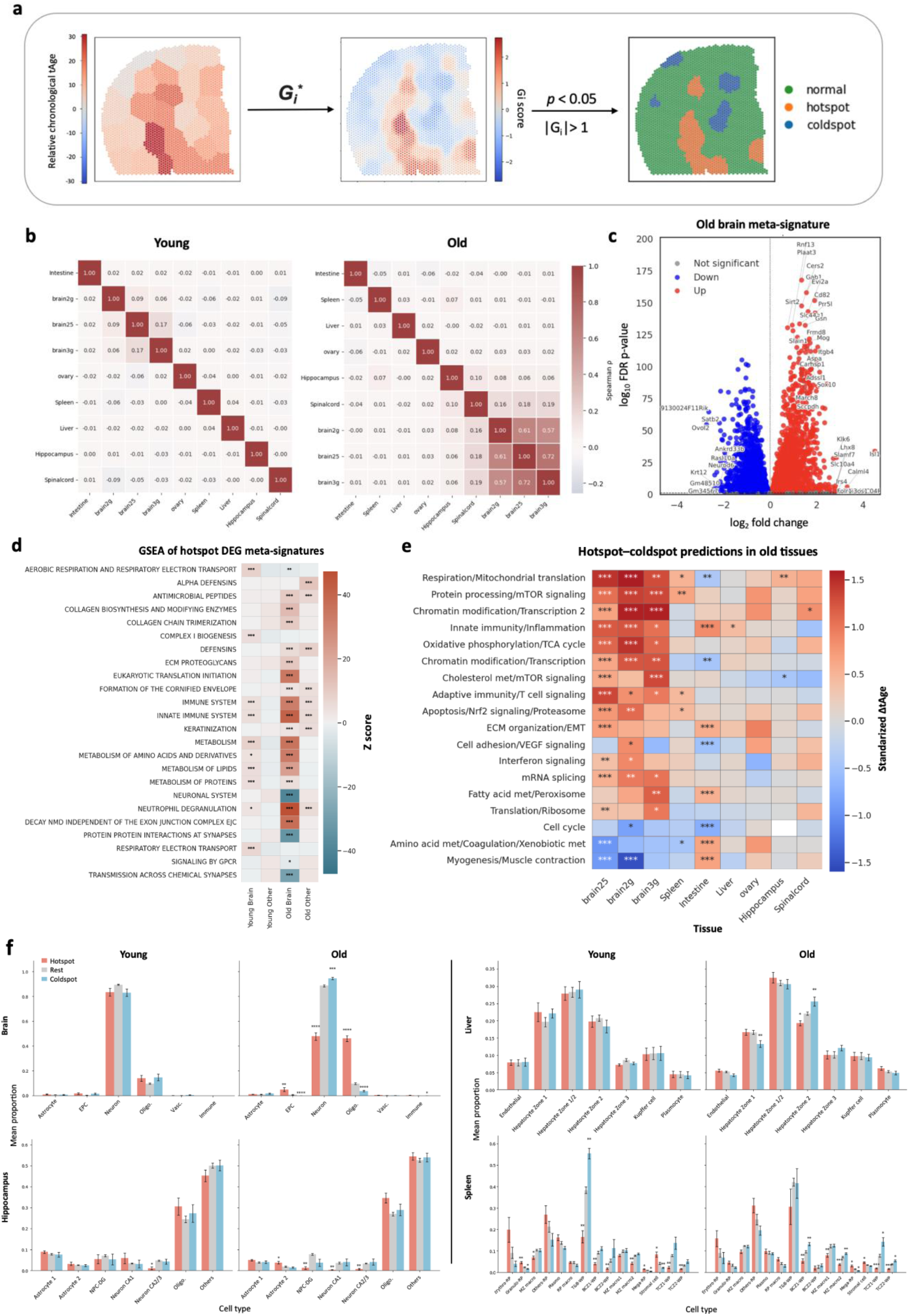
Signatures of regional biological age variability in natural aging. a. Identification of regional age acceleration. The G_i_* statistic was used to identify regions with significant increase (hotspots) or decrease (coldspots) in transcriptomic age. **b. Correlation of hotspot gene signatures.** Correlation between DEGs in hotspots across datasets. **c. Meta-signatures of aged hotspots in old brain.** Inverse variance weighting was used to draw gene meta-signatures across old brain datasets. **d. Enriched pathways of aged hotspots.** Self-contained gene set enrichment analysis was performed in young and old metasignatures of brain and other tissues. Top 9 pathways per sample are displayed. **e. Module clock signatures of hotspots in old tissues.** Standardized difference in tAge between hotspots and coldspots was calculated for each module clock, indicating mechanisms contributing to biological age difference. Mann-Whitney U test. **f. Cell type abundance in differentially aged regions.** Average cell type proportions in hotspots, coldspots, and normal regions were deconvoluted and calculated across tissues. Error bars are the SE (Mann-Whitney U test). Significance: *P* < 0.05 (*), *P* < 0.01 (**), *P* < 0.001 (***), *P <* 0.0001 (****); adjusted P-values shown.

Differential expression between hotspots and all other spots was performed per tissue, yielding a set of tissue-specific hotspot signatures. Correlations between these signatures across tissues (Fig. 6b) revealed that brain datasets clustered together, as expected, while other tissues shared weaker similarity. Importantly, hotspot signatures showed limited positive correlation across tissues in young animals, but became more correlated in old animals, suggesting that distinct organs develop convergent transcriptional programs of localized age acceleration with advancing age.

To integrate results across tissues, we computed metasignatures by pooling per-tissue hotspot fold changes using inverse-variance weighting (Fig. S5). In young tissues, hotspot signatures were weak and did not yield significant meta-signatures, consistent with largely stochastic variability. In contrast, old hotspots exhibited strong and shared metasignatures, especially in the brain (Fig. 6c), indicating that local age acceleration becomes more structured and reproducible with age.

Gene set enrichment analysis of these metasignatures (Fig. 6d) showed that, relative to coldspots, hotspots across tissues are enriched for immune and inflammatory pathways, including neutrophil degranulation and innate immune responses. In aged brain, hotspots also showed upregulation of lipid and amino acid metabolism, translation initiation and extracellular matrix synthesis, and downregulation of synaptic and oxidative phosphorylation pathways. Tissue-specific analyses (Fig. S6) corroborated a widespread upregulation of immune pathways in most tissues, with the exception of ovary (non-significant, possibly due to low sample size), and downregulation in spleen and intestine.

To quantify pathway-specific contributions to regional age differences, we applied module-specific transcriptomic clocks, previously trained on co-regulated gene modules enriched for distinct cellular processes^11^. These module clocks estimate how much each gene module contributes to the tAge difference between two groups—in this case, hotspots versus coldspots. In young tissues, few modules differed significantly between hotspots and coldspots (Fig. S7), in line with weak hotspot signatures. In old tissues, however, many modules showed significant contributions to hotspot–coldspot tAge differences, supporting pronounced local age acceleration (Fig. 6e). This effect was strongest in large, anatomically complex samples, with whole-brain sections displaying more spatial aging variability than smaller regions such as isolated hippocampus.

Across brain datasets, top module contributors to hotspot age acceleration included mitochondrial translation and respiration, mTOR signaling, chromatin modification, innate immunity and inflammation, and oxidative phosphorylation. Separate module-clock comparisons of hotspots and coldspots (Fig. S8) reiterated that energy metabolism and immune signatures contribute to hotspot age acceleration and indicated that chromatin modification modules contribute most to the lower biological age of coldspots, pointing to distinct molecular programs of local frailty and resilience.

Because regional tAge differences might be driven by cell-type composition, we used existing spot type annotations or annotated cell type identities for spots in ST samples (Fig. S9) and compared cell type proportions between regions across tissues (Fig. 6f). Cell type proportions in young tissues were similar across hotspots and coldspots, reinforcing the notion that early hotspots are not driven by large compositional shifts and showing that regional age acceleration is not driven solely by cell type identity.

In contrast, in old tissues, brain hotspots were enriched for oligodendrocytes and endothelial progenitor cells and depleted for neurons, consistent with elevated inflammation and accelerated aging in white matter regions^13,16^. At higher spatial resolution (Stereo-seq), hippocampal hotspots showed reduced abundance of all neuronal subtypes and increased astrocytes, a cell type known to become chronically inflamed with age^24^. In other tissues, composition differences were subtler but detectable—for instance, liver coldspots were enriched for zone 1 and 2 hepatocytes, and spleen hotspots showed reduced macrophage-related populations.

### Multispecies stAge captures age difference in human aging and disease

Finally, we extended stAge to human ST data using multispecies multi-tissue transcriptomic clocks^11^ (Fig. 7). In the human dentate gyrus dataset^25^ (Fig. 7a), stAge accurately distinguished four predefined age ranges. In human cortical samples^26^ (Fig. 7b), stAge separated the two age groups, similarly to in mice, outer cortical layers seemingly exhibited lower tAge than inner regions enriched for white matter, indicating a potential conserved spatial gradient of brain aging.

**Figure 7.**
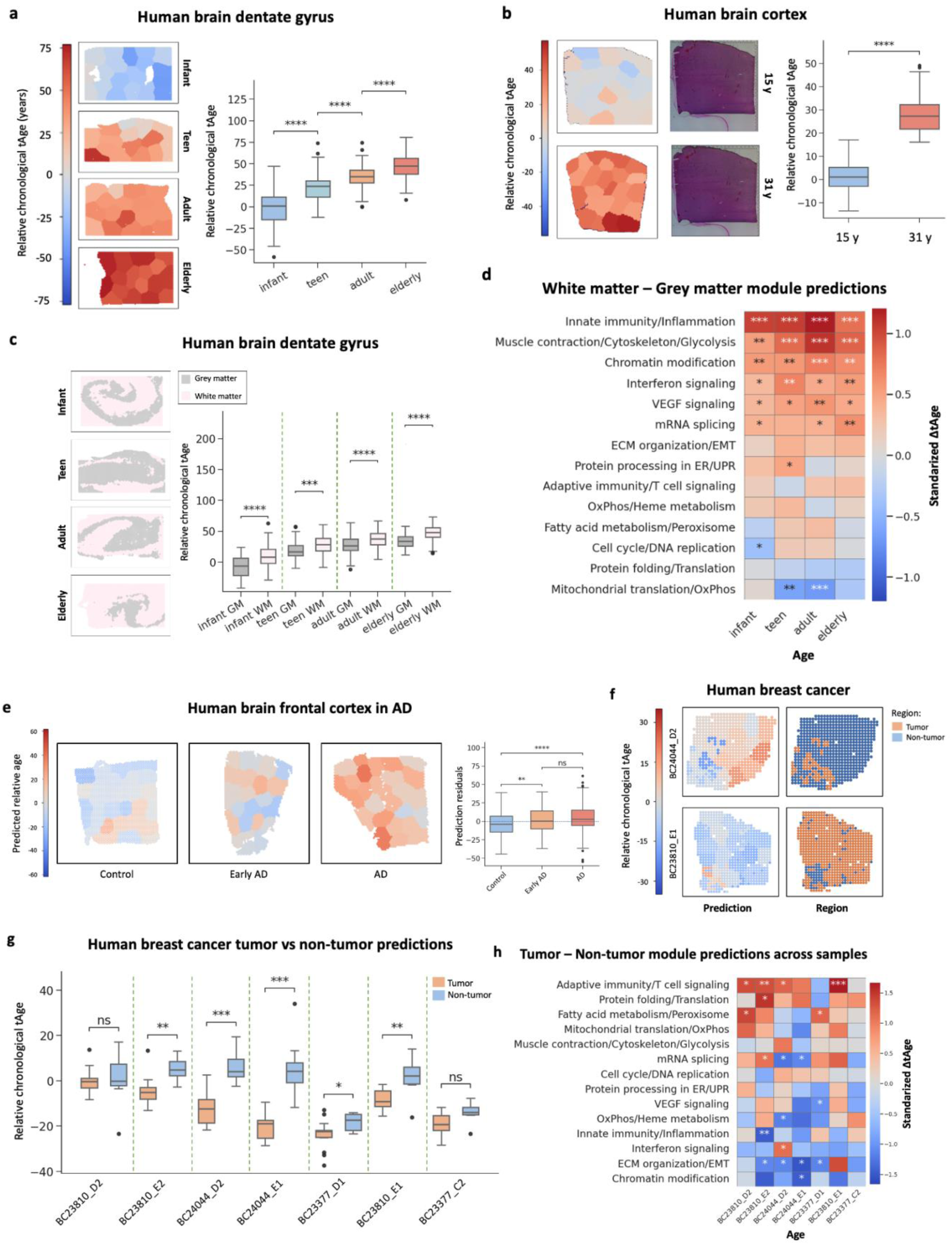
Multispecies stAge captures age difference in human aging and disease. a. Spatial age difference in the dentate gyrus (DG). Spatial and box plots of multispecies stAge relative age predictions (in years) in the dentate gyrus of infant (0-2 years old), teen (13–18 years old), adult (30–50 years old), and elderly (70+ years old) individuals (Zenodo ID 10126688). One sample in each age group were selected for the spatial plots. In box plots, all samples’ metaspot predictions are displayed for each time point. Mann-Whitney U test. **b. Spatial age difference in the cortex.** Spatial, H&E staining, and box plots of multispecies stAge relative age predictions in the cortex of 15- and 31-year-old individuals (GSE280570). One young and one old sample were selected for the spatial plots. In box plots, all samples’ metaspot predictions are displayed for each time point. Mann-Whitney U test. **c. White matter and grey matter transcriptomic age difference in the DG.** Marker genes were used to annotate WM and GM areas of the dentate gyrus, and WM-GM were compared at each age range. In every age range, white matter displayed higher biological age than grey matter. Mann-Whitney U test. **d. Module clock signatures of hotspots in old tissues.** Standardized difference in transcriptomic age between white matter and grey matter was calculated for each module clock, indicating mechanisms contributing to biological age difference. Mann-Whitney U test. **e. Spatial age difference in the human frontal cortex in Alzheimer’s disease (AD).** Human frontal cortex samples classified into control, early AD, and AD patients (GSE233208). Predictions were generated, and residuals of predicted relative age as a function of true age were collected for each group. Significantly higher residual deviation, reflecting age acceleration, can be observed in AD patients. Mann-Whitney U test. **f. Relative spatial age in breast cancer samples.** Spatial plots of multispecies stAge relative age predictions and tumor annotation in human breast cancer samples (Mendeley ID 10.17632/29ntw7sh4r.5). **g. Tumor and non-tumor regions transcriptomic age difference.** Tumor and non-tumor spot annotation was used to separate areas and predict their transcriptomic age separately. Within samples, tumor regions trend to lower biological age than non-tumor areas. Mann-Whitney U test. **h. Module clock signatures of tumor regions.** Standardized difference in transcriptomic age between tumor and non-tumor regions was calculated for each module clock, indicating mechanisms contributing to biological age difference. Mann-Whitney U test. Significance: *P* < 0.05 (*), *P* < 0.01 (**), *P* < 0.001 (***), *P <* 0.0001 (****); adjusted P-values shown.

To directly compare white matter (WM) and grey matter (GM), we annotated WM and GM regions in the human dentate gyrus and evaluated tAge across age ranges (Fig. 7c). In every age group, WM displayed significantly higher tAge than GM, confirming that the white-matter–associated aging gradient observed in mice is conserved in humans. Using multispecies module clocks, we then quantified pathway-specific contributions to the WM–GM age gap (Fig. 7d). Across all life stages, innate immunity and inflammation was the dominant contributor, suggesting that chronic neuroinflammatory processes are a persistent, spatially localized driver of WM age acceleration^27,28^. Cytoskeleton and glycolysis modules contributed most strongly in adult samples, consistent with accelerated decline in energy production in WM^29^. Chromatin modification, interferon signaling and VEGF signaling also contributed significantly across age ranges, highlighting additional pathways underpinning sustained WM–GM age differences.

stAge also captured age acceleration and deceleration in human disease. In human Alzheimer’s disease (AD) frontal cortex, stAge distinguished AD from non-AD samples and detected marked cortical age acceleration, mirroring observations in the 5xFAD mouse model (Fig. 7e). In human breast cancer ST data, tumor regions generally exhibited lower tAge than adjacent non-tumor areas (Fig. 7f,g), driven largely by changes in the ECM organization and EMT module, but with a substantial opposing contribution from immune pathways in many samples (Fig. 7h).

Taken together, these results show that stAge generalizes across tissues, interventions and species, including humans, and reveals conserved spatial and molecular mechanisms of aging: from white matter-centric neuroinflammation to tissue-specific immune, metabolic and chromatin programs that shape local biological age.

## DISCUSSION

We present stAge, a framework for quantifying and interpreting biological age in spatial transcriptomics data by integrating transcriptomic aging clocks with spatial neighborhood clustering. By applying an elastic net model to spatial metaspots defined by transcriptional similarity and physical proximity, we overcome noise limitations of single-spot prediction and enable robust spatial mapping of biological age across tissues.

stAge reliably distinguished young from old samples across diverse mouse tissues, including brain, liver, spleen, and testis, and uncovered regional age heterogeneity within tissues. In aged samples, we identified discrete hotspots of biological age acceleration, which became more prevalent and transcriptionally coherent with chronological age. These regions were enriched in innate immune signatures, including neutrophil activation and antimicrobial responses, consistent with chronic inflammation as a pattern of aging^30,31^.

In the brain, accelerated stAge regions aligned with white matter tracts and showed elevated expression of lipid metabolism and ECM remodeling genes, alongside suppression of synaptic and mitochondrial programs—transcriptomic changes associated with neuroinflammation and oligodendrocyte aging^32,33^. Module-specific clocks further identified mTOR signaling, oxidative phosphorylation, and chromatin remodeling as key contributors to spatial aging variation, supporting known systemic drivers of age-related decline^34^. Immune processes generally contribute to hotspot age acceleration, whereas chromatin-modification genes are the strongest contributors to the lower biological age of coldspots.

Cell type deconvolution revealed that aged hotspots were enriched in oligodendrocytes, endothelial progenitors, and astrocytes, and depleted in neurons. The enrichment of reactive astrocytes aligns with their known role in propagating neuroinflammatory signals during brain aging^35,36^. However, the lack of enrichment of a particular cell type in young tissue hotspots shows that cell type composition alone did not fully explain regional age acceleration, suggesting a role for both microenvironmental or cell-intrinsic regulatory changes.

Applying stAge to human cortex and dentate gyrus validated its cross-species generalizability, capturing conserved aging gradients such as older transcriptomic age in white matter compared to grey matter, largely driven by increased immune and metabolic aging signatures, which is consistent with known structural and cellular vulnerabilities during aging^27–29,37^.

While immune activation is a common hallmark of aging, we observed that hotspots in the intestine and spleen exhibited downregulation of immune-related pathways based on DEG enrichment. In the intestine, this likely reflects immune exhaustion, epithelial senescence, or loss of mucosal defense mechanisms, while in the spleen, hotspots showed suppression of adaptive immunity and T cell signaling. However, despite this apparent immune silencing, immune-related module clocks—targeting innate immunity in the intestine and adaptive immunity in the spleen—still contributed strongly to predicted age differences. This suggests that the absence of immune gene expression may itself act as an aging signal in regions undergoing structural or immunological decline. Notably, intestinal hotspots were also enriched in low-turnover compartments such as smooth muscle, whereas coldspots aligned with high-turnover epithelial zones. This turnover gradient may further confound differential expression and module analyses, as reduced regenerative capacity and baseline immune quiescence in low-turnover regions likely intersect with transcriptomic aging signatures^38,39^.

The predictive accuracy of stAge is dependent on the quality and tissue coverage of the training dataset. As the clock was trained on multi-tissue bulk RNA-seq, predictions in underrepresented or specialized tissues may be less reliable. Metaspot clustering is based on spatial and transcriptomic proximity but may not align perfectly with histological boundaries. Spatial transcriptomic technologies also vary widely in resolution and sensitivity, complicating cross-sample comparisons. While we observed consistent patterns of regional age acceleration, causality cannot be inferred; some changes may reflect reactive or compensatory processes. Finally, broader application to human tissues will require expanded spatial transcriptomic datasets, which remain limited in scope and resolution.

Together, these findings demonstrate that biological age is not only tissue-specific but also spatially patterned within tissues, with reproducible aging niches emerging in both mouse and human datasets, largely characterized by a shift in immune pathway regulation and chromatin modification. stAge enables mechanistic exploration of spatially localized aging processes and may inform spatially targeted interventions in aging and disease.

## METHODS

### Datasets

Both natural aging and intervention datasets were used for the study. Spatial transcriptomics data were obtained from published studies and the repositories NCBI GEO, STOmicsDB, GSA, Zenodo, and Mendeley with the following identifiers: GSE266933, GSE188257, GSE193107, GSE212903, GSE202322, GSE284202, GSE233208, GSE256397, GSE176092, GSE297119, GSE280570, GSE132042, GSE212336, STT0000039, PRJNA1043093, 10126688, 10.17632/29ntw7sh4r.5.

### Training and validation of transcriptomic aging clocks

Mouse and multi-species multi-tissue transcriptomic clocks of relative chronological age based on Elastic Net model were derived from Tyshkovskiy et al. (2024)^11^. To train the clock based on the meta-dataset encompassing both tissue samples and samples from single-cell RNAseq reflecting aging-related changes within individual cell types, we extended our original meta-dataset with Tabula Muris^40^ metacell data ^11^, using low-coverage (50,000 counts per metacell) and high-coverage (500,000 counts per metacell) thresholds for metacell aggregation. Gene expression was normalized by scaling or YuGene transformation, and hyperparameters were optimized through cross-validation, as described ^11^.

### Spatial metaspot clustering using SpatialGroup

To delineate spatially coherent transcriptional domains, we developed SpatialGroup, a method that clusters spatial transcriptomic spots into biologically and anatomically meaningful metaspots by integrating transcriptomic similarity and physical proximity. For each sample, a spatial neighbor graph was constructed using Squidpy’s sq.gr.spatial_neighbors, which defines adjacency based on a k-nearest neighbors strategy in spatial coordinates. The graph was then input as the adjacency matrix to Scanpy’s implementation of the Leiden community detection algorithm (sc.tl.leiden), ensuring that resulting clusters respect both spatial continuity and transcriptional structure.

In contrast, performing Leiden clustering on transcriptomic similarity alone (i.e., without spatial constraints) recapitulates expected cell type partitions but lacks anatomical coherence, highlighting the importance of spatial priors for metaspot definition.

The Leiden algorithm seeks to maximize modularity *Q* under a resolution-dependent objective function:

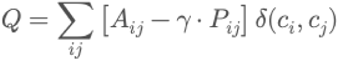

where:

- *A_ij_* is the observed weight between nodes *i* and *j*,
- *P_ij_* is the expected weight under a null model (e.g., configuration model),
- *δ(c_i_,c_j_)* = 1 if nodes *i* and *j* are in the same cluster, and 0 otherwise,
- *γ* is the resolution parameter, which controls the relative importance of within-cluster density.

Higher values of \gamma promote finer partitions by penalizing large clusters, thereby increasing the number of resulting metaspots.

### Regional annotation and clustering

Certain tissue samples with strongly distinguished regions had their spots annotated according to these regions. The intestine was separated based on spot type: high-turnover luminal regions (comprising stem and crypt cells, enterocytes, and goblet cells) and low-turnover peripheral regions (consisting of smooth muscle cells, plasmocytes, Paneth cells, and lymphocytes).

In the brain, the isocortex, hypothalamus, hippocampus, olfactory cortex, fiber tracts, thalamus, and striatum were annotated using curated sets of canonical marker genes as defined by Wang et al., (2025)^13^. For each dataset, expression values were normalized, log-transformed, and reduced to the top 2,000 highly variable genes, followed by PCA and graph-based clustering with the Leiden algorithm. To assign biological identities, we computed the mean cluster-level expression of each marker set, z-scored across clusters, and scored clusters by the average z-score of their respective regional markers. Each cluster was annotated with the region showing the highest score, with a confidence threshold of 0.5 to avoid spurious assignments. Clusters falling below this threshold were labeled as Unknown. This procedure yielded consistent automatic annotation of major brain regions across samples while preserving flexibility to update or refine marker catalogues in future analyses.

In other datasets, namely the skeletal muscle injury^20^ and AD model^22^, the original disease region annotation from the study was provided and respected. For region-specific predictions, the different regions were split into different objects, run through the pipeline, and joined back together after generating predictions.

### Optimal resolution search

The resolution parameter in the Leiden algorithm dictates the cluster size, increasing the pooled spots at higher values. To maximize resolution (γ) and look at tissues more in-depth but also account for the fact that tAge prediction performance differs across tissues and increases with more pooled gene coverage, a composite score (*S*) was developed to objectively determine the optimal resolution value for Leiden clustering in each dataset.

*S* is defined as:

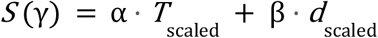

*S* is a function of Cohen’s d (*d*), measuring how well young and old samples are separated by magnitude, and the T statistic (*T*), measuring how statistically significant the separation between young and old samples is. Generally, higher resolution values generate more metaspots, which increase T, but smaller coverage per metaspot, which reduces tAge performance, thus decreasing *d*. *T* and *d* values are scaled after being computed so they account to make their magnitudes comparable. *α* and *β* are the weights assigned to each parameter, here set to 0.4 and 0.6, respectively – this assigns slightly higher importance to maximizing Cohen’s d and ensuring good separation between young and old samples.

After computing S for a given tissue at multiple resolutions, the highest resolution parameter value γ(*S*) within *S_max_* ± 10%.

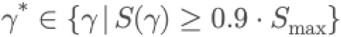

Tolerance of 10% was established to maximize resolutions with similar performance and obtain more granular but still reliable predictions.

### Sample processing and clock prediction

After metaspots (samples) are generated and their gene expression pooled, genes with less than 10 raw counts in at least 20% of samples were filtered out. Raw counts were added 1, log-transformed, and each sample was normalized separately with StandardScaler() – the same procedure followed to train the “Scaled” elastic net model. Additionally, samples were YuGene-transformed before running the “YuGene” elastic net.

The designated reference group, usually young samples, was subtracted to all sample groups to make gene expression relative. As shown before, relative age predictions improve the accuracy of the model. This results in the control groups prediction oscillating around 0 and all other predictions being relative to it. Missing genes were imputed with average precalculated values. After processing, samples’ gene expression was inputted into the elastic net models and predictions were generated for each of them.

### Lung infection and mouse AD model age acceleration quantification

Average transcriptomic age of metaspots was calculated for every day post infection (0, 3, 9) in young and aged mice. This value and the standard error of the mean were used to fit a mixed effects model of biological age acceleration with infection progression, for both YuGene and Scaled clock predictions.

### Hotspot and coldspot classification (Gi*) via Local Spatial Autocorrelation

To delineate spatial domains of transcriptomic age acceleration or deceleration, we employed the local Getis–Ord *G\** statistic, a spatial autocorrelation metric that quantifies the degree of local enrichment relative to global context. Specifically, for each spot i, the statistic is defined as:

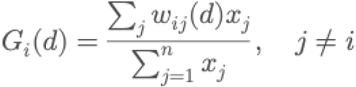

Where *x_j_*denotes the transcriptomic age at location *j*, and *w_ij_*(*d*) defines the spatial weight between *i* and its neighbors at distance *d*, constructed via a row-standardized k-nearest neighbor matrix with *k = 8*.

To assess statistical significance, we performed 999 Monte Carlo permutations of spot labels to generate empirical null distributions, computing standardized *G_i_\** z-scores and corresponding p-values. Multiple testing was controlled across all spots using the Benjamini–Hochberg FDR. Spots with FDR-adjusted *p* < 0.05 and *G \** > 1 were annotated as hotspots, indicating regions with significantly elevated biological age relative to their spatial context. Conversely, spots with *G\** < −1 and significant FDR were classified as coldspots, reflecting locally decelerated aging. All other spots were designated as non-significant.

### Differential expression analysis

Differential expression between hotspots and coldspots was assessed using a pseudobulk strategy. For each tissue dataset, spot-level counts were aggregated by hotspot vs coldspot within each sample. The resulting gene-by-sample pseudobulk count matrices were analyzed with DESeq2 in R.

The DESeq2 pipeline included dispersion estimation and negative binomial fitting. Hotspot vs coldspot contrasts were extracted and effect sizes were moderated using adaptive. Genes were annotated with log₂ fold-change, Wald test p-values, and Benjamini–Hochberg adjusted FDR values.

Standard errors for log-fold changes were further derived from two-sided p-values using the *scipy.stats.norm* distribution, enabling integration into downstream meta-signature analyses. Only tissues with at least two biological replicates per hotspot and coldspot group were retained for analysis.

### Correlation analysis

To evaluate consistency of hotspot-associated DEG signatures across tissues, we constructed ranked gene vectors from DESeq2 effect sizes. For each tissue, genes were ranked by log₂ fold-change (hotspot vs coldspot), and these vectors were concatenated into a tissue-by-gene matrix. Pairwise correlations between tissues were computed using Spearman’s ρ. The correlation matrix was converted to a distance metric (1 − ρ) and hierarchically clustered with average linkage. This analysis provided a comparative overview of tissue-specific and shared transcriptional programs distinguishing hotspots from coldspots.

### Gene signature meta-analysis

To integrate differential expression results across tissues, we applied a random-effects meta-analysis at the gene level. For each tissue-specific comparison of hotspots vs coldspots, log-fold changes and their standard errors were extracted. Genes missing standard errors were supplemented with values back-calculated from effect sizes and p-values.

A DerSimonian–Laird style estimator was implemented: within-tissue variances were used to compute fixed-effect weights, followed by estimation of between-tissue variance (τ²) and heterogeneity (I²). Pooled effect sizes, standard errors, z-scores, and Benjamini–Hochberg adjusted FDR values were obtained for each gene. This produced a set of standardized “meta-signatures” summarizing hotspot–coldspot contrasts across tissues.

### Gene set enrichment analysis

Meta-signatures were tested for functional enrichment using a self-contained gene set enrichment analysis (GSEA). For each gene set, standardized gene-level z-scores were combined with the Stouffer/Lipták method, with an optional variance inflation factor to account for average correlation between genes. This adjustment prevents inflation of test statistics due to coordinated expression within gene sets.

Gene sets were drawn from curated libraries (KEGG Hallmark and Reactome) and merged into a single dictionary. Pathways with fewer than 10 matched genes were excluded. Significance was assessed with two-sided tests, and FDR correction was applied across all tested terms. The resulting enrichment profiles highlighted biological pathways consistently associated with transcriptomic aging hotspots.

### Mouse module clocks

To dissect the biological underpinnings of transcriptomic aging at the tissue level, we utilized module-specific mouse and multi-species multi-tissue clocks trained to predict chronological age using subsets of individual clusters of co-regulated genes enriched for specific cellular pathways^11^, such as inflammation, chromatin regulation, oxidative phosphorylation, or mTOR signaling

Module-specific clocks were applied to spatial metaspots across tissues using the same pipeline as composite clocks. To evaluate the contribution of each module to aging differences, we compared standardized biological age estimates between young and old samples. For a given tissue *t* and module clock *m*, we computed the standardized effect size:

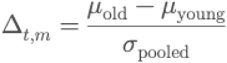

where:

- μ_old_, μ_young_ are the mean predicted ages in hotspot and coldspot samples, respectively,
- σ_pooled_ is the pooled standard deviation across both groups:

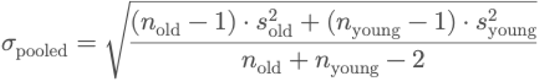

To simplify computation, we alternatively used the standard deviation of the concatenated vector of all predicted values in the tissue. Differences were then expressed as z-scored deltas. Statistical significance was assessed using Welch’s t-test on the standardized distributions.

Δ values and p-values were compiled across tissues and modules to generate summary heatmaps. Rows (modules) were ranked by their mean absolute Δ across tissues to highlight modules most consistently associated with age acceleration or deceleration.

### Cell type deconvolution

Cell type identities in the brain datasets were assigned to spatial transcriptomic profiles of spots using a supervised reference-based approach implemented in R. For each dataset, processed H5AD objects were converted to Seurat and harmonized with a curated single-cell reference containing high-confidence cell type annotations^35^.

Reference and query datasets were normalized using log-transformed counts following size factor estimation with *computeSumFactors*. Cell type prediction was performed with *SingleR*, which assigns each query profile to the most similar reference transcriptomic signature. The resulting labels were mapped back to the spatial coordinates, yielding spot-level cell type annotations consistent with the reference atlas.

### Cell type abundance calculation

Cell type composition was quantified within spatially defined hotspots, coldspots, and normal regions. For each sample, the relative abundance of each annotated cell type was calculated as the proportion of spots belonging to that type within a given region. Abundances were stratified by age group to enable comparison between young and old tissues.

Statistical differences in cell type representation were evaluated using nonparametric tests. Specifically, Mann–Whitney U tests were employed for within-age comparisons between hotspot or coldspot regions and their normal counterparts, while Welch’s t-tests were used to assess differences between age groups. Multiple hypothesis testing was controlled using the Benjamini–Hochberg procedure. This framework provided quantitative evidence for shifts in cellular composition underlying the spatial organization of transcriptomic aging signatures.

### Annotation of grey and white matter regions in human brain

Grey and white matter (GM/WM) regions were annotated in human cortical spatial transcriptomics sections using canonical marker panels. A curated list of oligodendrocytic and myelination-associated genes (MBP, PLP1, MOG, MAG, MAL, OPALIN, CLDN11, CNP, OLIG1, OLIG2, SOX10, TF) defined WM, while neuronal markers (SLC17A7, SLC17A6, SLC6A1, GAD1, GAD2, RBFOX3, SNAP25, SYT1, CAMK2A, MAP2) defined GM. Gene symbols were matched robustly across datasets by normalizing case, removing Ensembl version suffixes, and expanding aliases (e.g. NEUN→RBFOX3). Mean expression scores for each gene set were computed using Scanpy’s *tl.score_genes*, and the WM–GM expression difference (ΔWM–GM) was used to classify each spot. A two-component Gaussian mixture model identified an optimal threshold separating WM and GM populations. Spatial consistency was further refined by propagating labels across neighboring spots using Squidpy spatial connectivities and iterative smoothing (β = 0.8 over one neighborhood ring). The resulting annotation delineated cortical white- and grey-matter territories, coherent with histological tissue architecture, for downstream analyses.

### Multi-species module clocks

In parallel, we constructed gene module clocks based on the multi-species tAge model, a regularized regression clock trained on transcriptomic data from multiple organisms and tissues, also comprising ∼16,000 gene features. Functional modules were extracted and used to retrain separate elastic net clocks, using only the genes in each module. This allowed us to assess whether conserved aging mechanisms across species map to specific biological processes.

Predictions from these module clocks were applied to spatial metaspots and analyzed identically to the mouse module clocks. For each tissue and module, the standardized effect size between young and old samples Δ_t,m_ and its associated Welch p-value were computed and assembled into summary matrices. Rows were sorted by average |Δ| across tissues, and statistical annotations were added via p-value thresholds (e.g., p < 0.05, p < 0.01, p < 0.001).

### Human AD residual age analysis

To quantify biological age deviations independent of chronological age, we fitted an ordinary least squares (OLS) model of the form

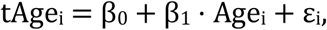

where tAge_i_is the relative transcriptomic age predicted for sample i, and Age_i_ is its true chronological age. Parameters β_0_ and β_1_ were estimated either across all samples or using only controls. The residuals (ε_i=_ tAge_i_ – β_0_ + β_1_ · Age_i_) were then used as a measure of transcriptomic age acceleration, representing the difference between predicted and expected age given the population trend.

### Breast cancer dataset

Small breast cancer samples with less than 600 observations (spots) were filtered out. Tumor and non-tumor annotation was provided by the authors. For each sample, module clocks predictions between tumor and non-tumor areas were used to derive contribution of each gene module to standardized tAge difference.

## DATA AVAILABILITY

This study did not generate original datasets. All spatial transcriptomics data analyzed were obtained from publicly available repositories or directly requested from original authors. The corresponding GEO accession numbers and dataset sources are listed in the Methods section. Any additional data supporting the findings of this study are available from the corresponding author upon reasonable request.

## ACKNOWLEDGEMENTS

Supported by NIH and Hevolution grants to VNG.

## SUPPLEMENTARY

**Figure S1.**
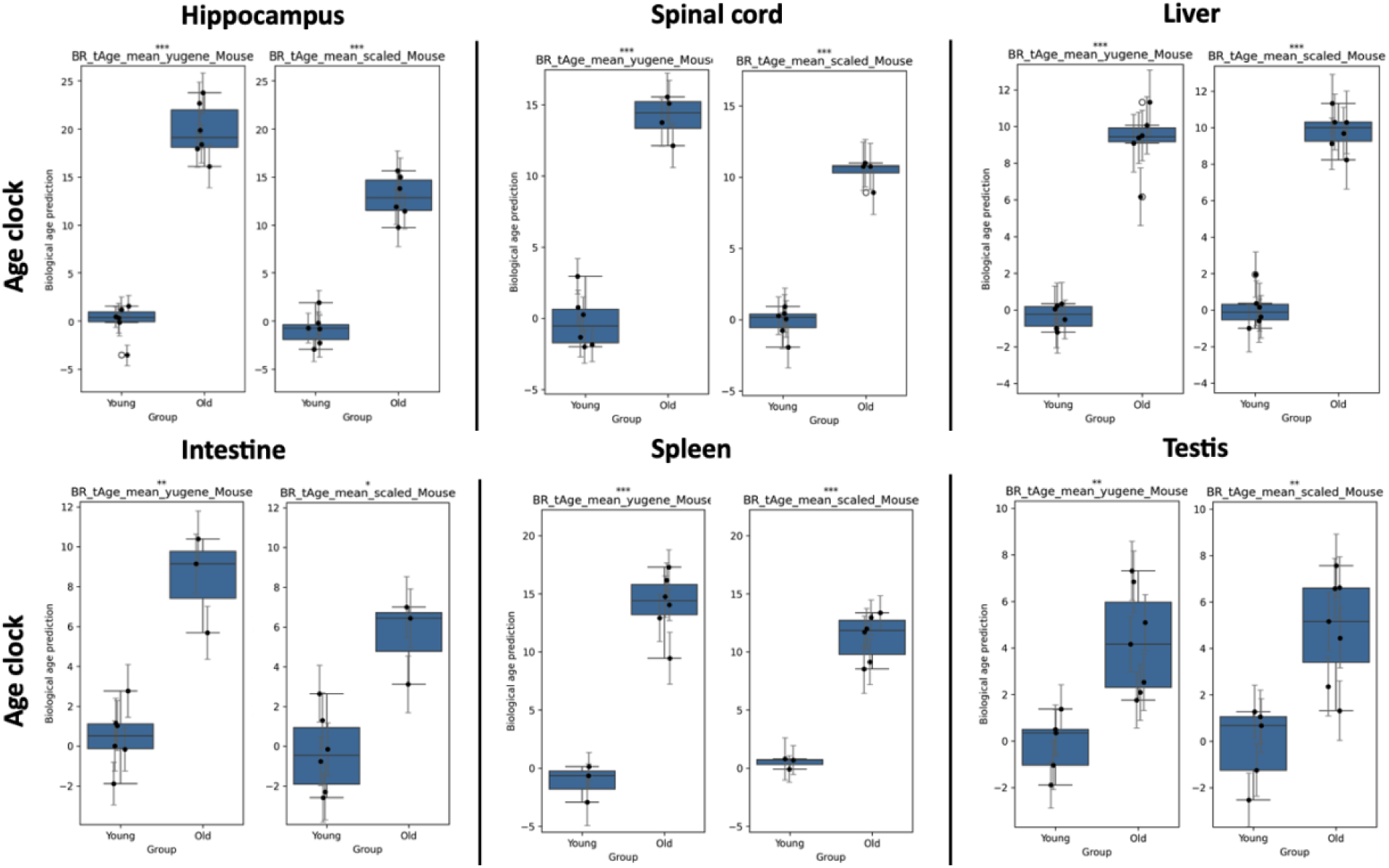
Pseudobulked Bayesian regression model predictions. For each gene, all counts in each spatial transcriptomics sample were summed. Pseudobulked transcriptomics samples were used to generate predictions with the Bayesian Regression tAge clocks trained on relative chronological age, both YuGene and Scaled versions. Box plots display prediction values and their standard error. Mixed effects modeling was used to characterize significance between young and old prediction difference.

**Figure S2.**
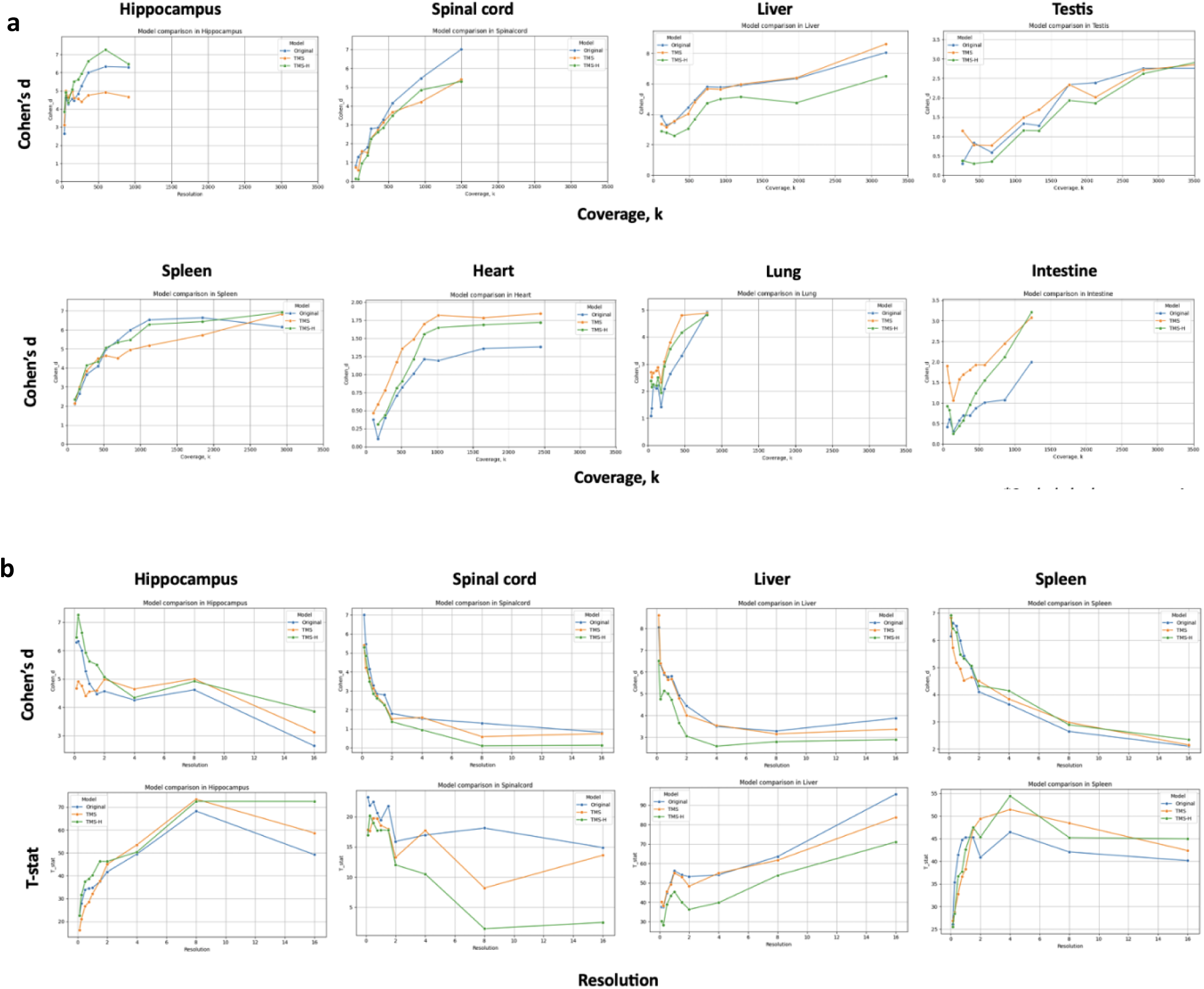
Performance of different stAge versions. a. Model performance comparison according to gene count coverage in multiple tissues. Metaspot clustering was done at different resolution values in different tissues, the average metaspot coverage for each resolution was calculated, and samples were used for predictions with three models: original tAge (Original), fine-tuned with low-coverage bootstrapped single-cell data (TMS), and fine-tuned with high-coverage bootstrapped single-cell data (TMS-H). Resolution and coverage are inversely proportionate. Due to differences in baseline expression across tissues, horizontal axes are set to the same dimension. All models’ Cohen’s d, reflecting their capacity to separate young and old samples, increases with higher coverage, but increased metaspot coverage reduces spatial resolution. **b. Model performance metrics according to resolution.** Metaspot clustering was done at different resolutions, predictions were obtained for each model, and Cohen’s d and T-stat was calculated between young and old samples, as a measure of the model’s separation capacity and significance. Overall, with higher resolution values (lower coverage per metaspot), Cohen’s d increases but T-stat decreases.

**Figure S3.**
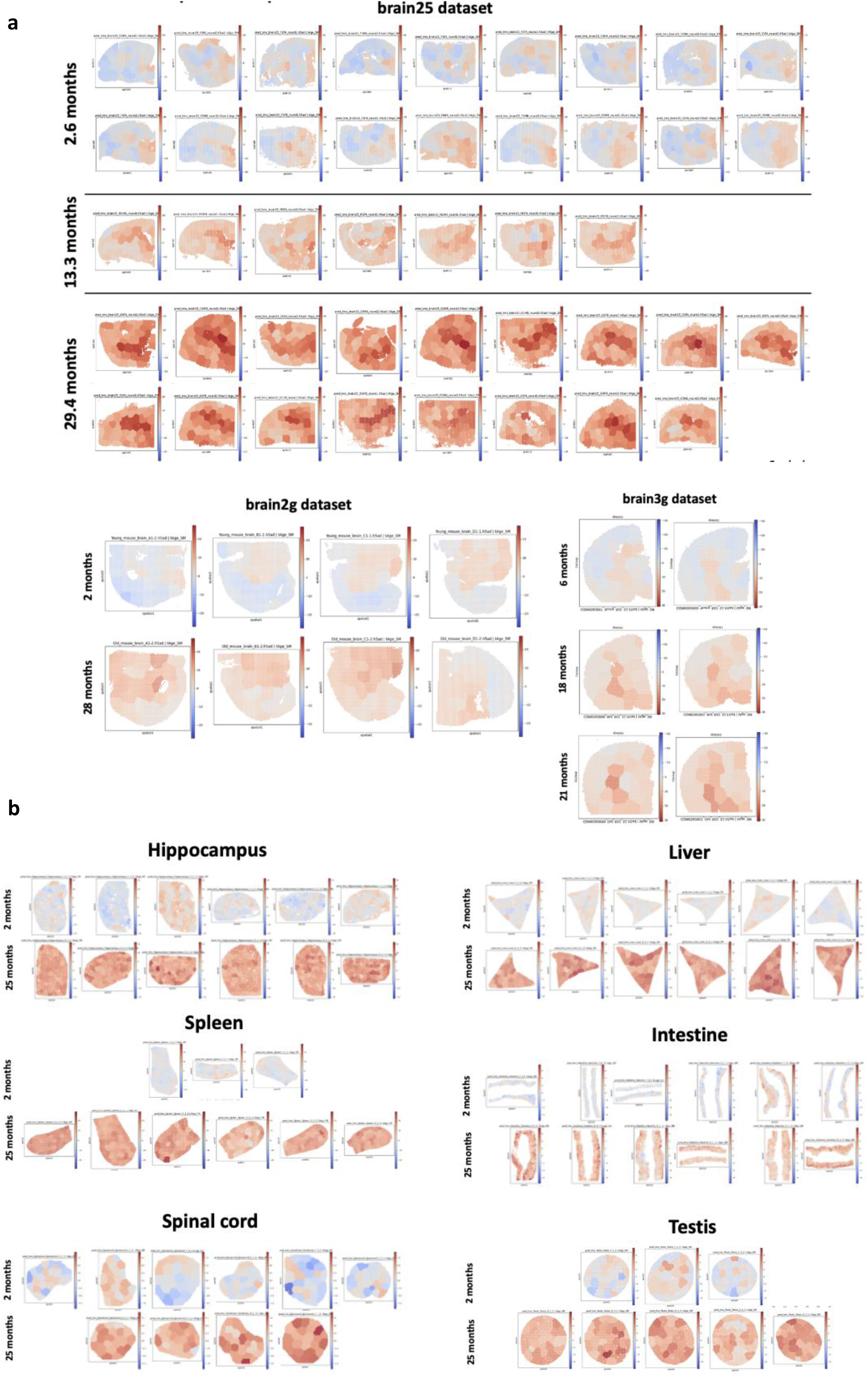
stAge age predictions of all samples using the scaled clock. a. Brain dataset predictions. Spatial plots showing predictions for young and old samples in three brain datasets: brain25 (GSE284202), brain2g (GSE193107), and brain3g (GSE212903). **b. Other dataset predictions.** Spatial plots showing predictions for young and old samples in the other natural aging datasets: Hippocampus, Spleen, Spinal cord, Liver, Intestine, Testis (STT0000039) and ovary (GSE188257).

**Figure S4.**
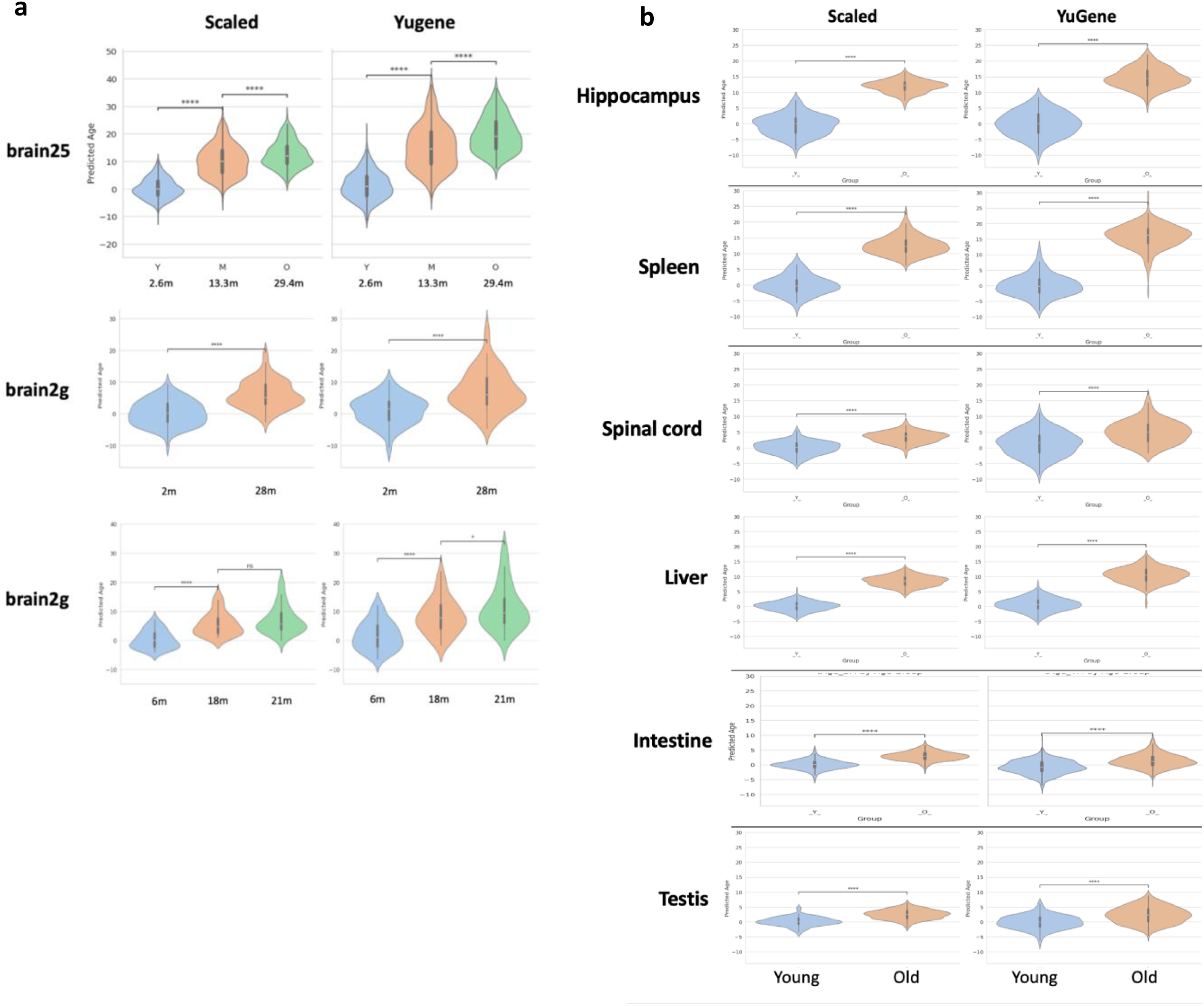
Scaled and YuGene clocks predictions. a. Brain dataset predictions. Violin plots showing metaspot age predictions of both Scaled and YuGene stAge versions for young, middle, and old samples in the three brain datasets: brain25 (GSE284202), brain2g (GSE193107), and brain3g (GSE212903). **b. Other dataset predictions.** Violin plots showing metaspot age predictions of both Scaled and YuGene stAge versions for young and old samples in the other natural aging datasets: Hippocampus, Spleen, Spinal cord, Liver, Intestine, Testis (STT0000039) and ovary (GSE188257). YuGene stAge predictions have generally higher variance. Mann-Whitney U test.

**Figure S5.**
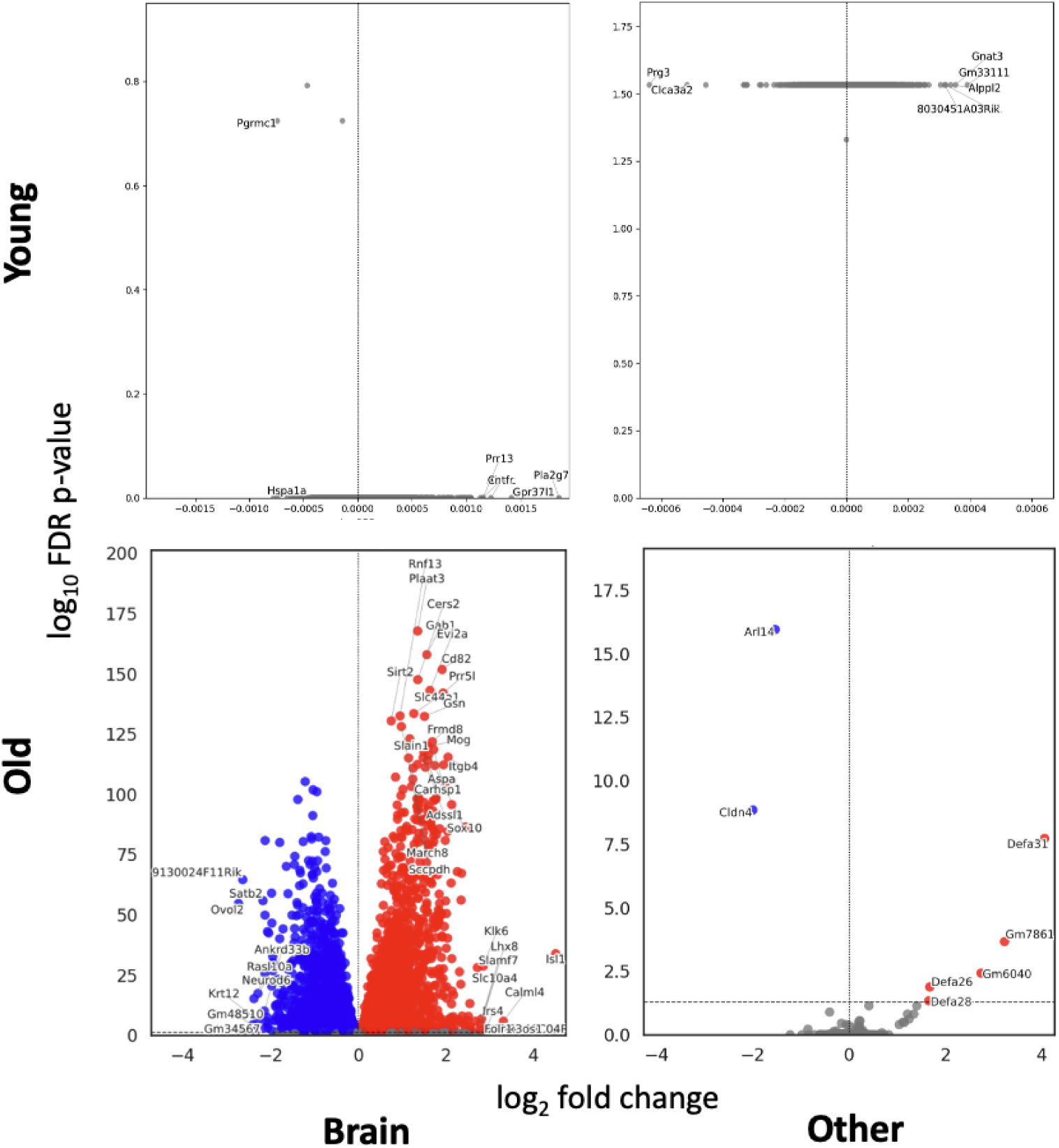
Meta-signature volcano plots. For each gene, hotspot-to-coldspot fold-change and adjusted p-value meta-signatures were calculated via inverse variance weighting. Meta-signatures were calculated for young brain datasets, other young datasets, old brain datasets, and other old datasets. Many more common significant genes were found across old tissues compared to young ones, especially across old brains, likely due to larger sample dimension and higher tissue regionality.

**Figure S6.**
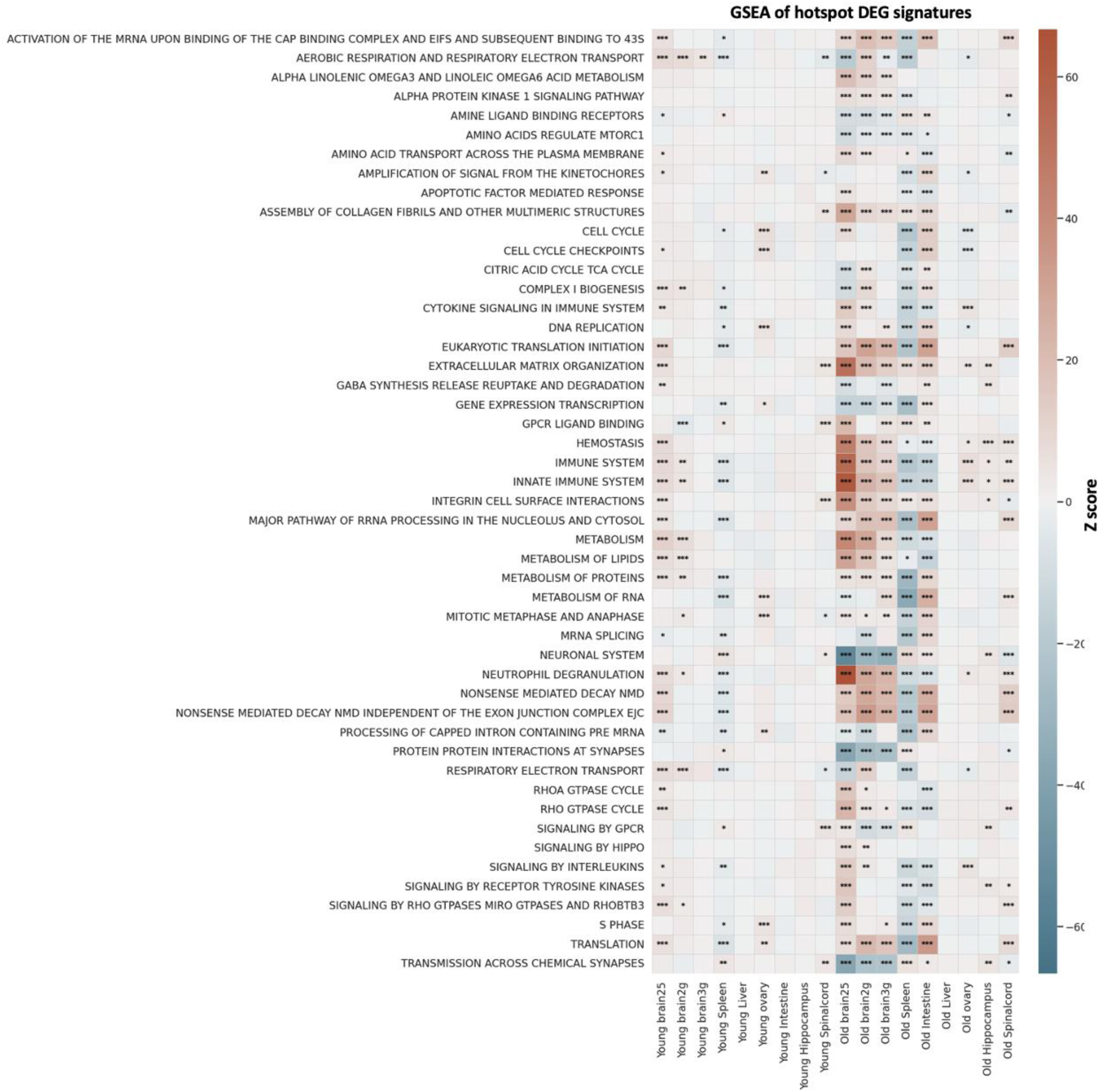
Enriched pathways of age-accelerated hotspots. Self-contained gene set enrichment analysis was performed in young and old signatures of different tissues. Top 5 pathways per sample are displayed.

**Figure S7.**
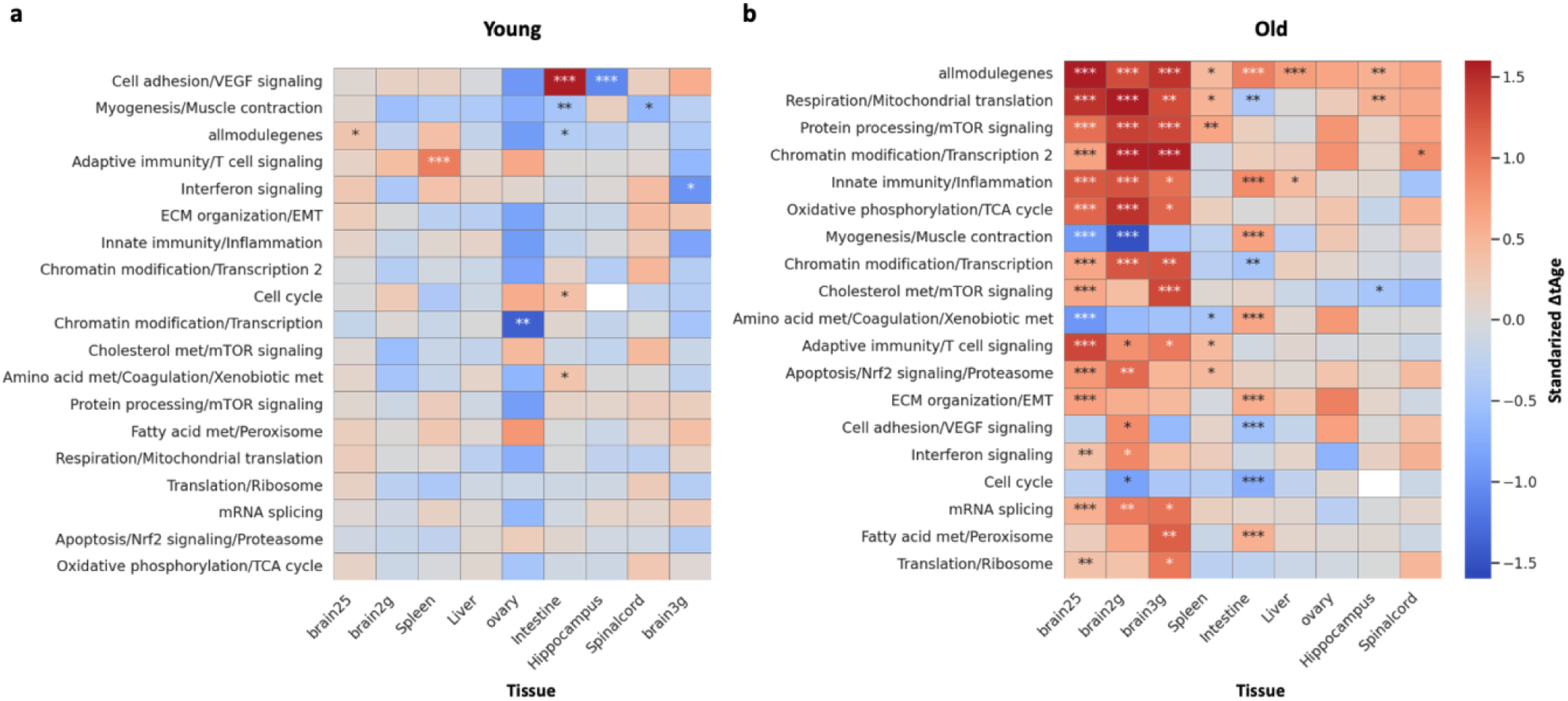
Module clock prediction hotspots vs coldspots in young (a) and old (b) tissues. Standardized difference in transcriptomic age between hotspots and coldspots was calculated for each module clock, indicating mechanisms contributing to biological age difference. Mann-Whitney U test.

**Figure S8.**
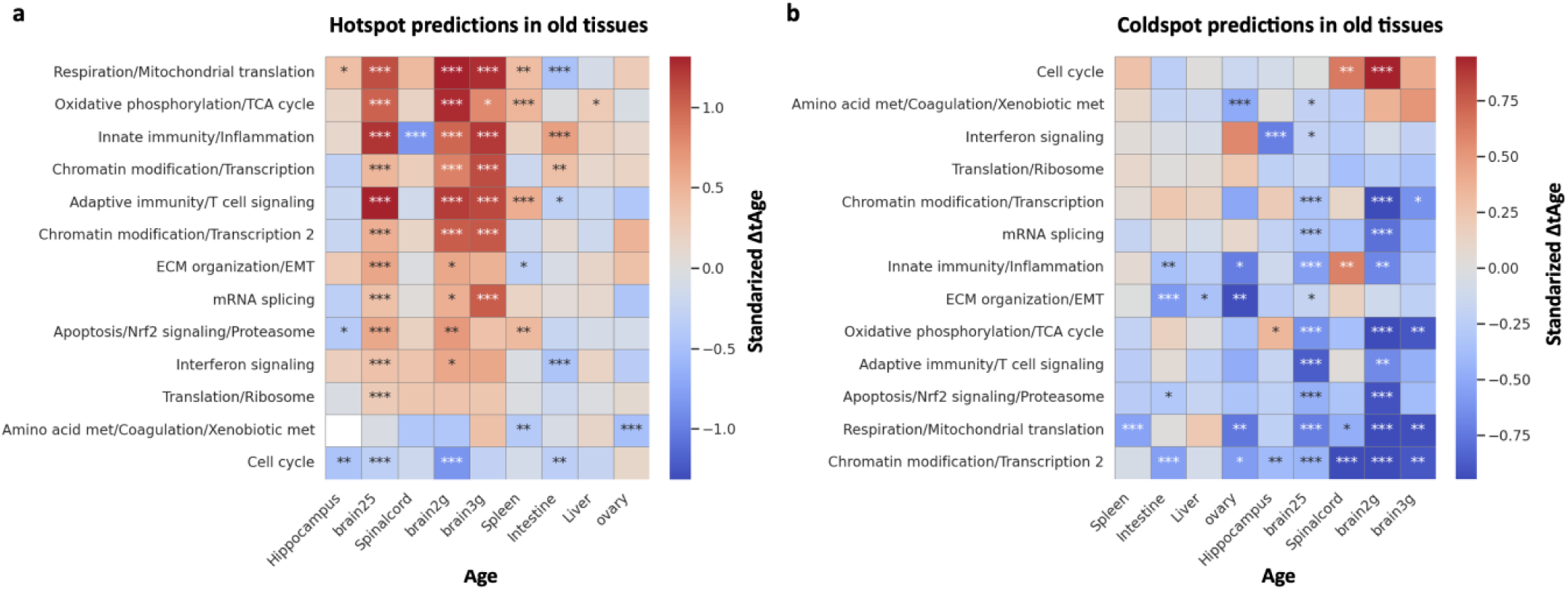
**a. Module clock prediction hotspots vs rest (normal and coldspot areas) in old tissues**. Standardized difference in transcriptomic age between hotspots and rest was calculated for each module clock, indicating mechanisms contributing to biological age difference. Mann-Whitney U test. **b. Module clock prediction coldspots vs rest (normal and hotspot areas) in old tissues**. Standardized difference in transcriptomic age between coldspots and rest was calculated for each module clock, indicating mechanisms contributing to biological age difference. Mann-Whitney U test.

**Figure S9.**
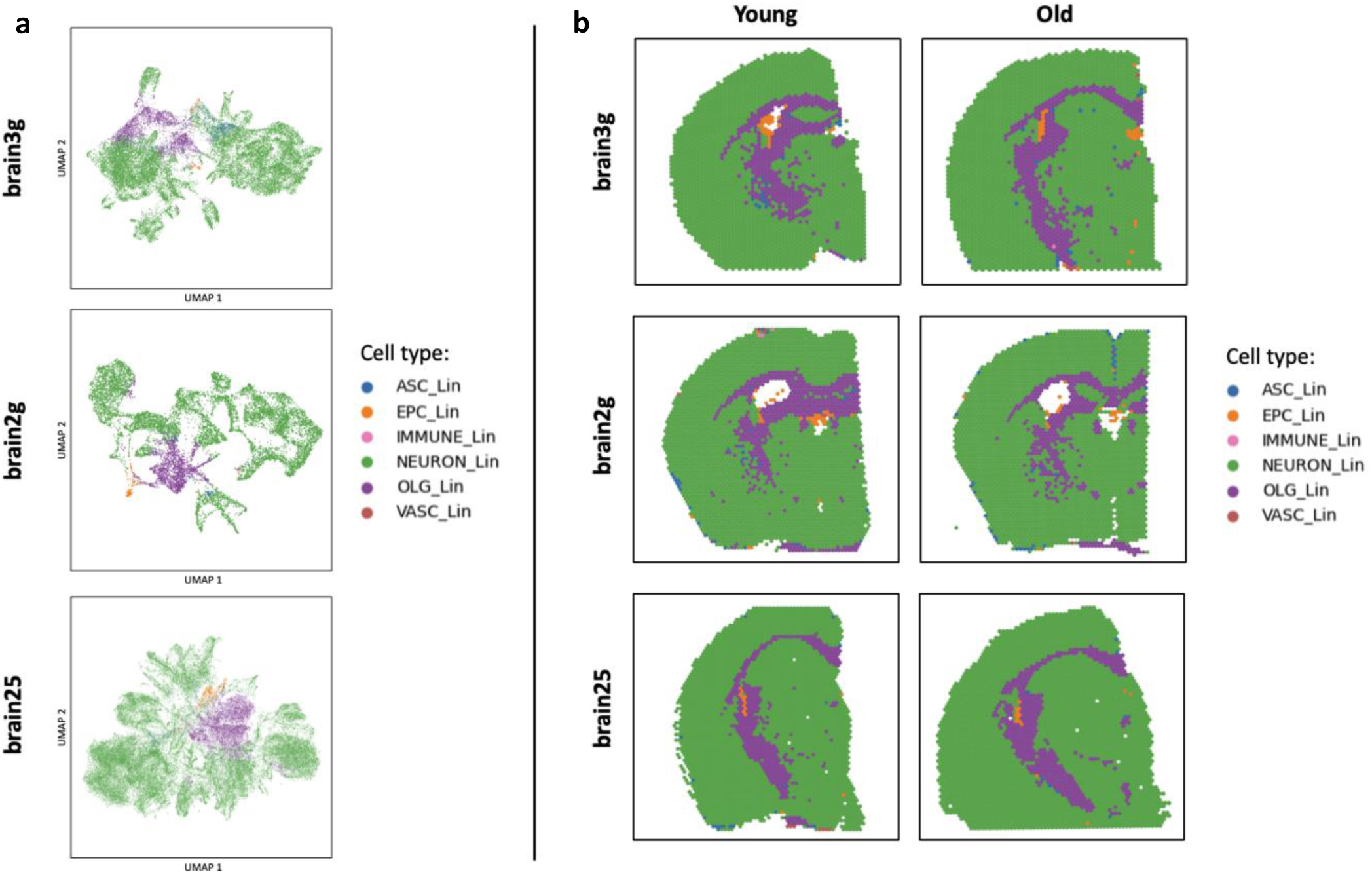
Cell type deconvolution. a. UMAP colored by cell-type annotation. Reference-based cell type deconvolution performed with singleR is shown to separate spots based on cell types in all three brain datasets: brain25 (GSE284202), brain2g (GSE193107), and brain3g (GSE212903). **b. Spatial plots colored by cell-type annotation.** Spatial plots show regional organization of cell types assigned to each spot via reference-based singleR cell type annotation in the three brain datasets.

## REFERENCES

1. Hou, Y. et al. Ageing as a risk factor for neurodegenerative disease. Nature Reviews Neurology vol. 15 565–581 Preprint at 10.1038/s41582-019-0244-7 (2019).

2. Tyshkovskiy, A. et al. Distinct longevity mechanisms across and within species and their association with aging. Cell 186, 2929–2949.e20 (2023).

3. Robeck, T. R. et al. Multi-species and multi-tissue methylation clocks for age estimation in toothed whales and dolphins. Commun Biol 4, 1–11 (2021).

4. Goeminne, L. J. E. et al. Plasma protein-based organ-specific aging and mortality models unveil diseases as accelerated aging of organismal systems. Cell Metab 37, 205–222.e6 (2025).

5. Oh, H. S. H. et al. Organ aging signatures in the plasma proteome track health and disease. Nature 624, 164–172 (2023).

6. Jin, K. et al. Brain-wide cell-type-specific transcriptomic signatures of healthy ageing in mice. Nature 638, 182–196 (2025).

7. Tian, L., Chen, F. & Macosko, E. Z. The expanding vistas of spatial transcriptomics. Nature Biotechnology vol. 41 773–782 Preprint at 10.1038/s41587-022-01448-2 (2023).

8. Stickels, R. R. et al. Highly sensitive spatial transcriptomics at near-cellular resolution with Slide-seqV2. Nat Biotechnol 39, 313–319 (2021).

9. Teschendorff, A. E. & Horvath, S. Epigenetic ageing clocks: statistical methods and emerging computational challenges. Nat Rev Genet 26, 350–368 (2025).

10. Sun, E. D. et al. Spatial transcriptomic clocks reveal cell proximity effects in brain ageing. Nature 638, 160–171 (2025).

11. Tyshkovskiy, A. et al. Transcriptomic Hallmarks of Mortality Reveal Universal and Specific Mechanisms of Aging, Chronic Disease, and Rejuvenation. Preprint at 10.1101/2024.07.04.601982 (2024).

12. Ma, S. et al. Spatial transcriptomic landscape unveils immunoglobin-associated senescence as a hallmark of aging. Cell 187, 7025–7044.e34 (2024).

13. Wang, L. et al. Spatial transcriptomics of the aging mouse brain reveals origins of inflammation in the white matter. Nat Commun 16, 3231 (2025).

14. Russ, J. E., Haywood, M. E., Lane, S. L., Schoolcraft, W. B. & Katz-Jaffe, M. G. Spatially resolved transcriptomic profiling of ovarian aging in mice. iScience 25, (2022).

15. Wu, C. et al. Spatially resolved transcriptome of the aging mouse brain. Aging Cell 23, (2024).

16. Hahn, O. et al. Atlas of the aging mouse brain reveals white matter as vulnerable foci. Cell 186, 4117–4133.e22 (2023).

17. Wang, Z. et al. A spatiotemporal molecular atlas of mouse spinal cord injury identifies a distinct astrocyte subpopulation and therapeutic potential of IGFBP2. Dev Cell 59, 2787–2803.e8 (2024).

18. Yamada, S. et al. Spatiotemporal transcriptome analysis reveals critical roles for mechano-sensing genes at the border zone in remodeling after myocardial infarction. Nature Cardiovascular Research 1, 1072–1083 (2022).

19. Wang, H., et al. Integrating spatial and single-cell transcriptomics to characterize mouse long bone fracture healing process. Commun Biol 8, (2025).

20. Walter, L. D. et al. Transcriptomic analysis of skeletal muscle regeneration across mouse lifespan identifies altered stem cell states. Nat Aging 4, 1862–1881 (2024).

21. Franzén, L. et al. Mapping spatially resolved transcriptomes in human and mouse pulmonary fibrosis. Nat Genet 56, 1725–1736 (2024).

22. Miyoshi, E. et al. Spatial and single-nucleus transcriptomic analysis of genetic and sporadic forms of Alzheimer’s disease. Nat Genet 56, 2704–2717 (2024).

23. Getis, A. & Ord, J. K. The Analysis of Spatial Association by Use of Distance Statistics. Geogr Anal 24, 189–206 (1992).

24. Gudkov, S. V. et al. An emerging role of astrocytes in aging/neuroinflammation and gut-brain axis with consequences on sleep and sleep disorders. Ageing Res Rev 83, 101775 (2023).

25. Ramnauth, A. D. et al. Spatiotemporal analysis of gene expression in the human dentate gyrus reveals age-associated changes in cellular maturation and neuroinflammation. Cell Rep 44, (2025).

26. Steyn, C. et al. A temporal cortex cell atlas highlights gene expression dynamics during human brain maturation. Nat Genet 56, 2718–2730 (2024).

27. Raj, D. et al. Increased white matter inflammation in aging- and alzheimer’s disease brain. Front Mol Neurosci 10, 271751 (2017).

28. Antignano, I., Liu, Y., Offermann, N. & Capasso, M. Aging microglia. Cellular and Molecular Life Sciences 2023 80:5 80, 1–18 (2023).

29. Goyal, M. S. et al. Loss of brain aerobic glycolysis in normal human aging. Cell Metab 26, 353 (2017).

30. Yousefzadeh, M. J. et al. An aged immune system drives senescence and ageing of solid organs. Nature 594, 100–105 (2021).

31. Franceschi, C., Garagnani, P., Vitale, G., Capri, M. & Salvioli, S. Inflammaging and ‘Garb-aging’. Trends in Endocrinology & Metabolism 28, 199–212 (2017).

32. Damoiseaux, J. S. et al. White matter tract integrity in aging and Alzheimer’s disease. Hum Brain Mapp 30, 1051–1059 (2009).

33. Clarke, L. E. et al. Normal aging induces A1-like astrocyte reactivity. Proc Natl Acad Sci U S A 115, E1896–E1905 (2018).

34. López-Otín, C., Blasco, M. A., Partridge, L., Serrano, M. & Kroemer, G. Hallmarks of aging: An expanding universe. Cell 186, 243–278 (2023).

35. Ximerakis, M. et al. Single-cell transcriptomic profiling of the aging mouse brain. Nat Neurosci 22, 1696–1708 (2019).

36. Liddelow, S. A. et al. Neurotoxic reactive astrocytes are induced by activated microglia. Nature 541, 481–487 (2017).

37. Jeffries, A. M. et al. Single-cell transcriptomic and genomic changes in the ageing human brain. Nature 646, 657–666 (2025).

38. Elmentaite, R. et al. Cells of the human intestinal tract mapped across space and time. Nature 597, 250–255 (2021).

39. Fulop, T. et al. Immunosenescence and inflamm-aging as two sides of the same coin: Friends or Foes? Front Immunol 8, 328099 (2018).

40. Almanzar, N. et al. A single-cell transcriptomic atlas characterizes ageing tissues in the mouse. Nature 583, 590–595 (2020).

